# Genome landscape and genetic architecture of recombination in domestic goats (*Capra Hircus*)

**DOI:** 10.1101/2025.05.15.654186

**Authors:** Alice Etourneau, Rachel Rupp, Bertrand Servin

**Author notes:** Corresponding Authors, AE; BS. E-mail addresses: RR.

## Abstract

**Background:** Recombination is a fundamental biological process, both in participating to the creation of viable gametes and as a driver of genetic diversity. Characterising recombination is therefore of strong interest in breeding populations. In this study, we used ∼50K genotyped data and pedigree from two French populations (Alpine and Saanen) of domestic goats (*Capra hircus*) to build sex-specific recombination maps, and to explore the genetic basis of two recombination phenotypes: genome-wide recombination rate (GRR) and intra-chromosomal shuffling.

**Results:** Sex-specific recombination maps showed higher recombination in males than females for both breeds (Alpine autosomal map size = 35.1M in males and 30.5M in females; Saanen map size = 34.0M in males and 29.0M in females). Heterochiasmy is particularly notable on small chromosomes, and at both ends of the chromosomes. Yet, no difference in shuffling has been found between populations. Genetic parameters on recombination phenotypes could only be estimated in males, due to lack of data in females. Both phenotypes are significantly heritable (h²=0.12 (0.03) for GRR and h²=0.034 (0.015) for shuffling, when pooling breeds). GWAS on male GRR identified several significant loci, including *RNF212*, *RNF212B* and *SSH1*, the last one being a novel locus for this phenotype. Correlation of SNP effects between breeds is low for both recombination phenotypes (0.25 for GRR and 0.04 for shuffling), indicating different genetic determinants in the two breeds.

**Conclusions:** This study contributes to the understanding of recombination evolution in ruminants, both between breeds and species.

## Background

Recombination is a fundamental biological process in eukaryotes. It is necessary for the proper segregation of chromosomes during meiosis [1] and is also a great driver of genetic diversity by creating new combinations of alleles and avoiding the accumulation of deleterious mutations over time [2,3]. Studying recombination can therefore be important both to understand functional reproduction in a species, and to study evolutionary drivers of genetic diversity.

Of the different approaches to study recombination, pedigree-based studies, as opposed to population studies for instance, have several advantages: first they provide a direct measure of meiotic recombination, essentially unaffected by evolutionary pressures, through the transmission of gametes in the pedigree and second, they allow to assign crossovers to specific individuals and therefore estimate the genetic basis of the recombination process. Pioneered in humans [4,5], the increased availability of cheap genotyping tools has led to a large number of recombination studies, especially in livestock. Indeed, the implementation of genotyping routines within genomic selection programs put at the disposal of researchers both the tools and the data to study recombination in pedigrees. As such, multiple studies were conducted in cattle [6–11], but also in sheep [12], chicken [13], pigs [14–17] and Atlantic salmon [18]. Apart from livestock, recombination in several natural populations with available pedigrees have also been studied: Soay sheep [19], red deer [20,21] and wild house sparrow [22].

This approach of characterising recombination through the use of genotypes and pedigree data allows to estimate recombination rates and build recombination maps of a species. Results have shown that heterochiasmy, or sex-differences in recombination, is widespread in many clades, but its sign varies: in most species like pigs [14–16], humans [4], deer [20], chicken [13] or Atlantic salmon [18], females recombine more than males; while in other species like cattle [8–11] or sheep [19], males show higher recombination than females. [22] and [23] illustrate this inconsistent heterochiasmy in birds and fishes, respectively.

This high diversity in recombination features shows the relevance of studying the recombination process itself. One way to study this process is through genetics, which allows to estimate heritability and find associated QTLs for phenotypes such as recombination rate and, more recently, intra-chromosomal shuffling [24]. The genetic control of recombination rate, despite showing common basis between species, was also shown to be very diverse, even between several populations or the two sexes of the same species. Focusing only on ruminants, heritability for phenotypes of recombination intensity varies between 0.04 in Norwegian Red cattle [11] and 0.22 in Dutch Holstein-Friesian cattle [6]. But variations in heritability can be observed between species and, most notably, sexes within a species [9,12,19,21]. Identified QTL for these phenotypes can also vary between populations and species. Cattle [6,8–11], sheep [12,19] and red deer [21] all show a signal in the region of the genes RNF212B/HEI10/REC8/REC114, four genes in very close proximity to each other in these three species. In cattle [6,9] and sheep [12,19], associations with the *RNF212* gene have also been found. Yet despite these similarities, genes like MSH4 or PRDM9 have only been detected in cattle among ruminants, although this is possibly due to the increased statistical power in this species. When studying genetic control within sexes and breeds separately in a species, the differences are even starker. In sheep, [19] found a QTL at RNF212 only in Soay females, and not in males. Yet [12] actually found a QTL for this same gene in males from another population, the Lacaune. In cattle, [8] identified thirteen loci associated with recombination intensity, and only three of them were shared between sexes. In [9]’s bovine population, four of their ten detected QTLs are sex-specific. And once again, [11] found sex differences in Norwegian red dairy cattle.

In this context of strong genetic diversity in very related groups, we present a new study on another ruminant species: the goat. In goats, the last genetic map was built with a linkage methodology in 1998 [25]. At the time, its primary goal was still to understand the organisation of genes between one another, and the map was never updated since. Genome-wide recombination rate was estimated through the use of cytogenetics techniques by counting MLH1 foci [26,27] but never confirmed using other methods.

In 2010, the first reference genome assembly was built for the domestic goat [28] and in 2013, an international goat genome consortium was created to acquire more genomic data and investigate genetic variations in the species. This lead to the development of a 50k SNP chip [29] and later on, the creation of a sequence-level SNP panel of more than a thousand goats from all over the world [30].

In France, a goat breeding program for two breeds, Alpine and Saanen, has been in place since the 1960s, its main goal being to improve milk production for the purpose of cheese-making with the progressive inclusion of health and udder morphology characteristics. Since 2014 and the availability of the 50k SNP chip, selection candidates are now systematically genotyped. Furthermore, despite the fact that the breeding program is male-oriented, females have started to be genotyped as well in the last few years, both as part of the breeding program and within experimental programs. Therefore, we took advantage of genotyping and pedigree data available in the two Alpine and Saanen breeds to study recombination in this ruminant species using a pedigree-based approach, for the first time to our knowledge. In an opportunity to examine heterochiasmy and breed differences in this new species, we built updated, sex-specific recombination maps, developing novel methodologies to quantitatively measure sex differences in recombination patterns and take into account uncertainty about crossover detection. In addition, we also studied the genetic basis of two recombination phenotypes in both available breeds, by estimating genetic parameters and performing genome-wide association studies.

## Methods

### Study population and genotype data

In this work, we exploited pedigree and genotyping data of 7,588 goats born between 2008 and 2019. They amount to 3,169 males and 4,419 females from both the Alpine (5,051 samples) and Saanen (2,537 samples) breeds. The animals were genotyped with versions 1 and 2 of the Illumina Inc. GoatSNP50 chip. Only autosomal SNPs and markers common to both arrays were kept, resulting in 53,347 SNPs.

### Quality control of physical marker ordering

Errors in the physical ordering of markers can lead to detect spurious recombination events. Recombination studies therefore require a robust marker ordering. Here, this ordering was built from two independent physical maps of the SNPs: the reference genome assembly ARS1 and the Radiation Hybrid (RH) map of [31]. We decided to keep only SNP markers that have a consistent ordering on both physical maps. This was done by identifying, for each chromosome, the longest increasing subsequence (LIS)[32] in the mapping of the RH order to the assembly order. This requires knowing the order of the SNP markers on both maps. For the reference genome ordering, genomic positions of the 53,347 v1 autosomal markers of the GoatSNP50 Illumina chip were obtained from the ARS1 assembly as described on the International Goat Genome website (http://www.goatgenome.org/projects.html). For the RH map, additional analysis was required as it was built using SNP array of different species (ovine and bovine SNPs herein called external SNPs). We mapped the external SNPs to the ARS1 reference genome and then projected the position of each caprine SNP on the RH map from the positions of its two flanking external SNPs. For goat markers located at the extremities of the chromosome, which had only one flanking external SNP and could not be projected on the RH map, we assumed the reference genome order was correct (but see below for further data quality control steps). **Figure S1** (**See Additional File 1**) illustrates this quality control step on two chromosomes. After this step, 54,013 markers were kept.

Furthermore, an initial detection of crossovers on the genome (using methods detailed below) revealed 7 genomic regions (123 SNP markers in total) that showed extremely high recombination rates (See example on chromosome 1 in **Additional File 1, Figure S2**). Detailed analysis of the synteny conservation with the human genome in these regions showed that they corresponded systematically to conserved synteny breakpoints. As these events were most probably some remaining unidentified assembly errors, markers involved were discarded in downstream analysis. For a list of these markers, see **Additional File 2, Table S1**.

### Genotype quality control

Standard data cleaning procedures were applied using PLINK 1.9 [33]. Animals and SNPs with call-rate below 99% were removed (**See Additional File 1, Figure S3**). Monomorphic SNPs in the dataset were removed. No MAF filter was applied at this step as even a small MAF may be informative for crossover detection in families polymorphic for these markers. Markers with strong departures from Hardy-Weinberg genotype proportions (*p* < 10^−7^) within each breed were removed. Note that families with a sire having more than 50 offspring were not used to compute the Hardy-Weinberg equilibrium (HWE) test as fathers carrying rare alleles in such designs can have a great impact on testing for HWE.

Mendelian errors were tested using the *mendel* command of the Yapp software version 0.3 [34]. It evaluated the number of incoherent markers between parent and offspring and automatically corrected the kinship relationships in the pedigree which were beyond the threshold fixed by the software.

After these quality control steps, 46,081 markers and 7,457 animals, including 4,520 Alpine and 2,937 Saanen, were left.

### Data phasing and crossover detection

Genotype phasing and crossover detection were done using Yapp version 0.4. First, the *phase* command was used in order to phase genotypes. In order to reconstruct the paternally and maternally inherited haplotypes of each individual (i.e. to infer the phase of heterozygous loci), Yapp uses a set of discrete steps. First, using all available genotypes, it identifies the unambiguous transmissions from parents to offspring, *e.g.* the alleles transmitted by a homozygous parent or received by a homozygous offspring are easily phased. This leads to a set of partially resolved phases. Second, for each parent in the pedigree, from oldest to newest, it improves the inference of parental phases by combining all partially resolved transmitted haplotypes using the algorithm of [35]. Third, after this first pass in the pedigree, the transmitted haplotypes from parent to offspring are improved using a Hidden Markov Model along the whole chromosome similar to the approach of [36] and segregation indicators are inferred. Then, the *recomb* command was used among a total of 6,434 meioses to detect crossovers. A crossover in the parent meiosis is defined as a change in phase observed in the offspring, on the inherited chromosome from the parent. Crossovers are detected between two heterozygous SNPs and can only be detected between the first and the last phased, heterozygous SNPs.

Meioses and crossovers were quality-controlled to ensure that only truly informative data was retained. All meioses with a parent or offspring phased at less than 90% of markers were removed. Furthermore, meioses with a parent that itself had none of its parents genotyped and only one offspring, were removed, as it is impossible to detect crossovers in such a setting.

Distance between two crossovers on the same chromosome was calculated for every meiosis. As crossovers are detected within an interval, distance between crossovers was calculated as the distance between the closest interval boundaries. All double-crossovers closer than 5 Mb from one another were considered false-positives due to phasing errors and both were removed (**See Additional File 1, Figure S4**). After filtering, the dataset contained 6,204 meioses and 204,727 crossovers.

### Recombination maps

Recombination maps were constructed by estimating recombination rates in non-overlapping intervals of one megabase (Mb) along the genome. The methodology for estimating recombination rates was adapted from [12]. For a given genomic interval *i* and a meiosis *m*, the count of the number of crossovers *Y*_*mi*_ is modelled using a Poisson distribution whose parameter (measured in Morgans) depends on the recombination rate *c*_*i*_ (in centiMorgans per Mb, cM/Mb) assumed constant in the interval and across meioses, and the length of the genomic interval *l*_*mi*_ (in Mb) that can depend on the meiosis (see below).

Then, the expected number of crossovers in an interval is 0.01*c*_*i*_*l*_*mi*_ and *Y*_*mi*_ is modelled as:

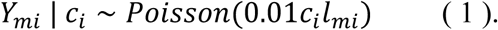

As in [12], we define a prior on *c*_*i*_ as:

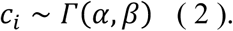

As in [12], we used an empirical Bayes approach to set *a* and *β* from the genome-wide distribution of Maximum Likelihood Estimates of recombination rates (*α* = 3.02 and *β* = 2.32, see **Additional File 1, Figure S5**). Given the conjugacy of Poisson and Gamma distributions, the posterior distribution for *c*_*i*_ is then:

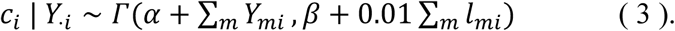

Stemming from the Bayesian model (3), the expected value for the Gamma law was used to estimate the chromosomic recombination rate:

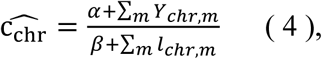

With *α* and *β* the prior parameters and *m* a meiosis. *c*_*chr*_ was estimated for both breeds and sexes, and a model explaining *c*_*chr*_ by the log of the chromosome size was fitted for all populations.

#### Meiosis Informativity

In the equations above, the length of the interval is allowed to depend on the interval and the meiosis considered. This is to account for the variation in the informativity of the data for the detection of crossovers. First, because samples are genotyped on a 50k SNP chip, no phasing information is available for the tips of chromosome before and after the first and the last SNP of the chromosome, which makes detecting crossovers impossible in these regions. Second, for a given meiosis, only SNPs that are heterozygous and phased in the parent are informative for crossover detection. Hence for a particular meiosis, this can further reduce the portion of the genome where crossovers can be detected (**Fig. 1**).

**Figure 1:**
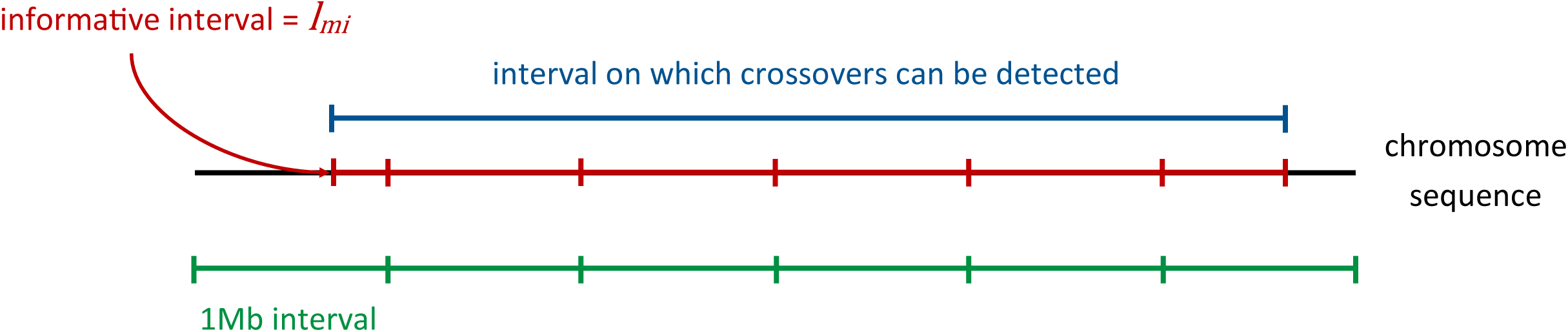
Illustration of the notion of interval informativity.

#### Accounting for uncertainty in crossover locations

Crossovers are not determined at a specific position but resolved within genome regions that vary in size depending on the parent and offspring genotypes. The average length of these intervals in our data is 1,4Mb with a 95% range of 0.1Mb to 10.2Mb (**See Additional File 1, Figure S6**). To account for this uncertainty in the position of crossovers, we used a Monte Carlo approach as in [12]: we first sampled crossover positions uniformly in the detection intervals and then the recombination rate from its posterior (Equation 4). This was done 100 times, providing for each interval 100 samples from the posterior distribution of recombination rate. Let us name *E*(*c*_*i*_|*Y*_⋅*ik*_) = *b*_*ik*_ the expected value of posterior *k* for genomic interval *i*. The expected value of all combined posteriors for interval *i* is therefore:

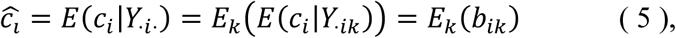

And according to posterior variance law:

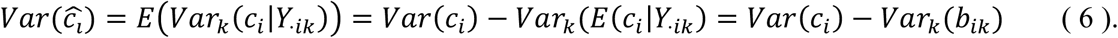

Thus, recombination rate *c* for genomic interval *i* was estimated as the mean of the 100 posteriors’ expected values.

#### Assessing sex differences in local recombination rates

The Bayesian model described above is interesting in that it allows to account naturally for the uncertainties in the data and provides estimates of recombination rates even in regions with little information. Using this approach, we could in principle evaluate the sex differences in recombination rates from the posterior samples obtained by applying the model separately in both sexes. However, when interrogating the difference in local recombination rates between sexes, the model does not allow to separate the effect of differences in rates and those of differences in uncertainty. Therefore, the two sexes could be deemed to have different rates due to the amount of data in each sex being unbalanced (for example female estimates are typically more influenced by the prior than male estimates). To deal with this issue, we derived a permutation scheme that can separate both effects.

Our measure of heterochiasmy in an interval *i* is:

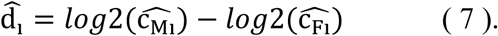

That is, the log-ratio of male to female recombination (in base 2 for easier interpretation).

To account for the difference in data informativity between sexes, we permuted the sex of the parents 1,000 times to obtain a sample of **d**_**i**_ under the null of no difference between sexes. As permutations do not change the number of male and female meioses, they allow to account for difference in informativity between sexes. The resulting distribution of **d**_**i**_ in the permutation samples were very close to normally distributed (**See Additional File 1, Figure S7**), so in each interval the null distribution of **d**_**i**_ can be described by a normal distribution with parameters set at the sample mean and variance. This in turn allows to derive p-values for a difference between sexes in each interval. Finally, to account for multiple testing across intervals, the False Discovery Rate (FDR) approach of [37] using the R “qvalue” package was applied on the p-values and the significance threshold set at 0.05.

### Recombination phenotypes

The crossover dataset described above offers the opportunity to study the individual variation in some aspects of recombination (i.e. recombination phenotypes) and interrogate their genetic architectures. However, because we had very little data in females this was only possible to do in males. Two phenotypes were studied: the individual Genome-wide Recombination Rate (GRR) and the intra-chromosomal shuffling.

The intensity of recombination in a meiosis can be measured by summing the crossover count on all autosomes (Autosomal crossover count (ACC)). However, as explained above, the informativity of marker data varies between meioses and can influence the value of ACC. To take this into account, rather than the ACC as a measure of recombination intensity we use the genome-wide recombination rate (GRR), defined as the ACC divided by the informative portion of the genome (in Mb).

The other phenotype we studied is a measure of the efficacy of a meiosis to create new allelic combinations in the offspring: the rate of intra-chromosomal shuffling *r̄*_*intra*_ [24]. It is defined as the probability that two randomly chosen loci uncouple their alleles during meiosis and is calculated as:

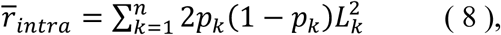

Where for chromosome *k*, *p*_*k*_ is the proportion of SNPs inherited from one parent of the focal animal (or grandparent of the offspring), *L*_*k*_ is the chromosome’s proportion in the autosomal genome, and *n* is the number of autosomes.

### Heritabilities

Heritabilities of the GRR and *r̄*_*intra*_ phenotypes were estimated using all males with a calculated phenotype, i.e. 520 parents accounting for 5,707 meioses (**Table 1** for detail).

**Table 1:**
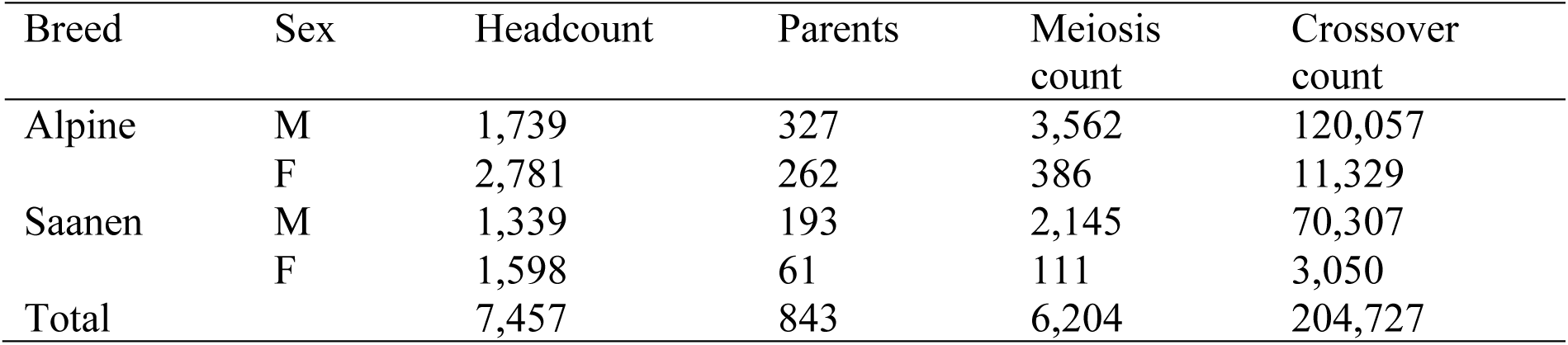
Summary table of data after all genotype and crossover quality control steps.

| Breed | Sex | Headcount | Parents | Meiosis count | Crossover count |
| --- | --- | --- | --- | --- | --- |
| Alpine | M | 1,739 | 327 | 3,562 | 120,057 |
|  | F | 2,781 | 262 | 386 | 11,329 |
| Saanen | M | 1,339 | 193 | 2,145 | 70,307 |
|  | F | 1,598 | 61 | 111 | 3,050 |
| Total |  | 7,457 | 843 | 6,204 | 204,727 |

To estimate heritability, variance components were estimated by restricted maximum likelihood (REML) using the Wombat software [38]. An animal model including a permanent environment effect was fitted:

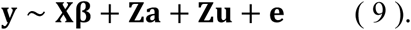

**y** is the vector of observations (GRR or *r̄*_*intra*_); **β** is the vector of fixed effects, **a** is the vector of direct animal (additive) genetic effects; **u** is the vector of animal permanent environment effect; **e** is the vector of residual effects; 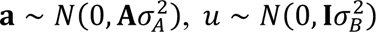 and **X**, **Z** are the incidence matrixes. The pedigree matrix **A** was computed on five generations of the genealogy. The model was applied to all breeds separately or to the entire population at once. A fixed effect for breed was therefore added where necessary.

Once the variance components had been estimated, values were incorporated into the model as fixed values to obtain the Estimated Breeding Values (EBV) for each phenotype. Due to the absence of differences in estimated heritability between breeds (see Results), EBVs were estimated by pooling samples from both breeds. Note that although we did not fit a model for female recombination phenotypes, EBVs can still be predicted for females as they are connected to males in the pedigree.

### Genome-wide association studies

#### Genotype Imputation

As an imputation reference panel, we used the whole-genome sequence VarGoats data presented in [30] and retrieved from https://www.goatgenome.org/home.html. This data consisted of 1,372 goats out of which 252 European samples closely related to our study samples were used (from the Alpine, Fossés, Lorraine, Poitevine, Provençale, Pyrénées, Rove, Savoie and Saanen breeds).

SNP filtering was then applied on the resulting VCF based on several criteria, using bcftools (http://www.htslib.org/doc/1.1/bcftools.html). First, we removed SNPs having discordant reference alleles between the SNP array manifest and the resequencing data, and SNPs showing strictly less than two counts of the minor allele across all selected samples or with a proportion of missing genotypes greater than 10%. Then, SNPs were filtered based on the distribution of the following GATK 3.6 [39] variant-calling quality parameters: DP, FS, VQSLOD (**See Additional File 1, Figure S8**). The chosen threshold for SNP removal were DP < 10,000; DP > 25,000; FS > 50; VQSLOD < 10 and VQSLOD > 25. The resulting number of SNPs remaining was 22,741,231 out of which 43,358 SNPs are present on the V1 GoatSNP50 Illumina chip. This corresponds to removing 5.6% of the chip SNPs.

The Beagle4 software [40] was used to impute the few missing genotypes of the sequence panel and correct the called genotypes by using the genotype likelihood (*gl*) method, a modelscale of 1.5 [41], an effective population size of 1000 and the recombination maps estimated using a pedigree-based approach *(see above)*. To assess the quality of the resulting panel genotypes, we leveraged information on 91 animals for which both sequencing and SNP genotyping data were available: we computed R² and genotype concordance before and after imputation. Both metrics were highly improved by the imputation procedure: mean R2 increased from 0.928 to 0.990 and mean genotype concordance from 0.974 to 0.997. (**See Additional File 1, Figures S9** & **S10**). Phasing of the reference panel was done using ShapeIt4 [42]. As some animals from the reference panel were previously phased using pedigree information, their phased haplotypes for the SNP chip markers were used as scaffold reference for phasing with ShapeIt4. The resulting VCF file, phased and imputed, was used as reference panel for imputation.

Shapeit4 was also run on the phased VCF of 7,457 samples produced by Yapp from the SNP array data to solve the few unphased genotypes and impute the few remaining missing genotypes. The resulting fully phased 50k chip data was imputed to sequence using the Impute5 software [43]. Recombination maps were provided to increase power and *N*_*e*_ was set at 1000. Impute5 provides a measure of confidence for all imputed markers (imputation R2) which was used in later steps to filter SNPs *(see below)*.

#### Statistical Analysis

GWAS were performed separately in each breed using the predicted EBV for male GRR and *r̄*_*intra*_ as phenotypes: 589 samples were used in Alpine and 254 in Saanen (Table 1). SNPs with MAF<5% and imputation R2<0.3 were removed from the analysis leaving N=15,988,092 and 15,734,503 SNPs in Alpine and Saanen respectively. A univariate linear mixed model was used for testing SNP association with the phenotype using the Gemma software [44], with the “leave one chromosome out” (LOCO) strategy (*i.e.* the genetic relationship matrix (GRM) used to analyse a chromosome is computed on all other chromosomes except this one).

Significance was assessed using the ash method [45] implemented in the “ashr” R package, by calculating s-values for each SNP. In the same way as for q-values [37], thresholding s-values at level α gives a set of significant features so that a proportion of α is expected to be false-positives. While [37]’s FDR gives indication on the confidence that effects are different from zero, [45]’s False Sign Rate (FSR) gives indication on the confidence of the *sign* of effects. As this method takes into account the *β* estimates as well as their standard errors, the resulting s-values can induce a reranking of SNPs compared to methods only taking p-values into account. All SNPs with s-value<0.01 were declared significant and SNPs with s-value<0.05 were declared suggestive.

Correlations of SNP effects between breeds were measured by the correlation of β estimates for SNPs that were significant in at least one of the two breeds, and using standard errors as weights.

#### Annotation of GWAS loci

A GWAS locus was defined from pairwise Linkage Disequilibrium (R2) between all significant SNPs on a given chromosome. R2 calculation was done using the PLINK 1.9 software. Significant SNPs were considered to belong to the same peak if their R2 was equal or superior to 0.2. For the lead SNP of each peak, a high LD area was defined as follows: an area whose boundaries correspond to the SNP furthest away in R2>0.6 from the lead variant, on both sides of the lead variant. We aimed at evaluating the evidence for association of all genes present in Ensembl (gff3 release 111). As this annotation did not include two major genes associated with the recombination process, *RNF212* and *PRDM9*, we manually added them to the annotation file after blasting their bovine gene sequence on ARS1 (**See Additional File 3, Tables S2 & S3**).

For all genes, we evaluated the following criteria: if the gene is within 2Mb of the lead variant; if it is the closest gene to the right, to the left or both; if it is the closest gene to another significant SNP of the peak and if it belongs within the “R2>0.6” area of the peak. Based on this we calculated a “candidate score”, ranging from 0 to 5, as the number of criteria satisfied by the gene.

## Results

### Recombination landscape

After quality control, genotype data on 7,457 animals led to the detection of 204,727 crossovers in 6,204 meioses from 843 parents. As a large portion of the dataset comes from a dairy selection program, most genotyped parents were males. As males also typically have more genotyped offspring than females, there are 10 times more male crossovers than female ones in the recombination data (Table 1). The software used to detect crossovers provides the information on the parts of the genome that are informative for the detection of crossovers (See Methods). Overall, all meioses had more than 94% of the genome informative for crossovers (**See Additional File 1, Figure S11**). Females had slightly more informative meioses than males (mean=0.99% vs 0.98%). Indeed, although females are usually less well connected in the pedigree than males, the ones that passed our quality control criteria for meioses (**see Methods**) tended to be better connected than the average male.

### Chromosome-wide rates

A linear model was used to test differences in genetic map size between populations. Breed and sex parameters were found significant (sex effect p-value = 10^−61^; breed effect p-value = 10^−13^; interaction p-value = 0.52). The size of the genetic map was estimated at 35.1M for Alpine males, 34.0M for Saanen males, 30.5M for Alpine females and 29.0M for Saanen females (**See Additional File 1, Figure S12**). Therefore, the genetic map is slightly shorter in Saanen than in Alpine and much shorter in females than males.

The mean recombination rate of each chromosome was estimated with the Bayesian model, and a linear model depending on the log size of the chromosomes was fit on the estimates (**Fig. 2a**). Again, a significant difference between breeds was found (p-value = 0.01) but, as the effect is very small, the figure shows breed-averaged estimates to avoid over-plotting. A significant difference between sexes is also observed (p-value ≈ 10^−4D^): males overall have a higher chromosomic recombination rate than females, and the recombination rate is greater on small chromosomes than big chromosomes for both sexes. Overall, the chromosomic recombination rate has a log relation with the chromosome size. It is observed that the difference in recombination between the sexes is greater on small chromosomes than on big chromosomes, on which recombination rates are almost alike.

**Figure 2:**
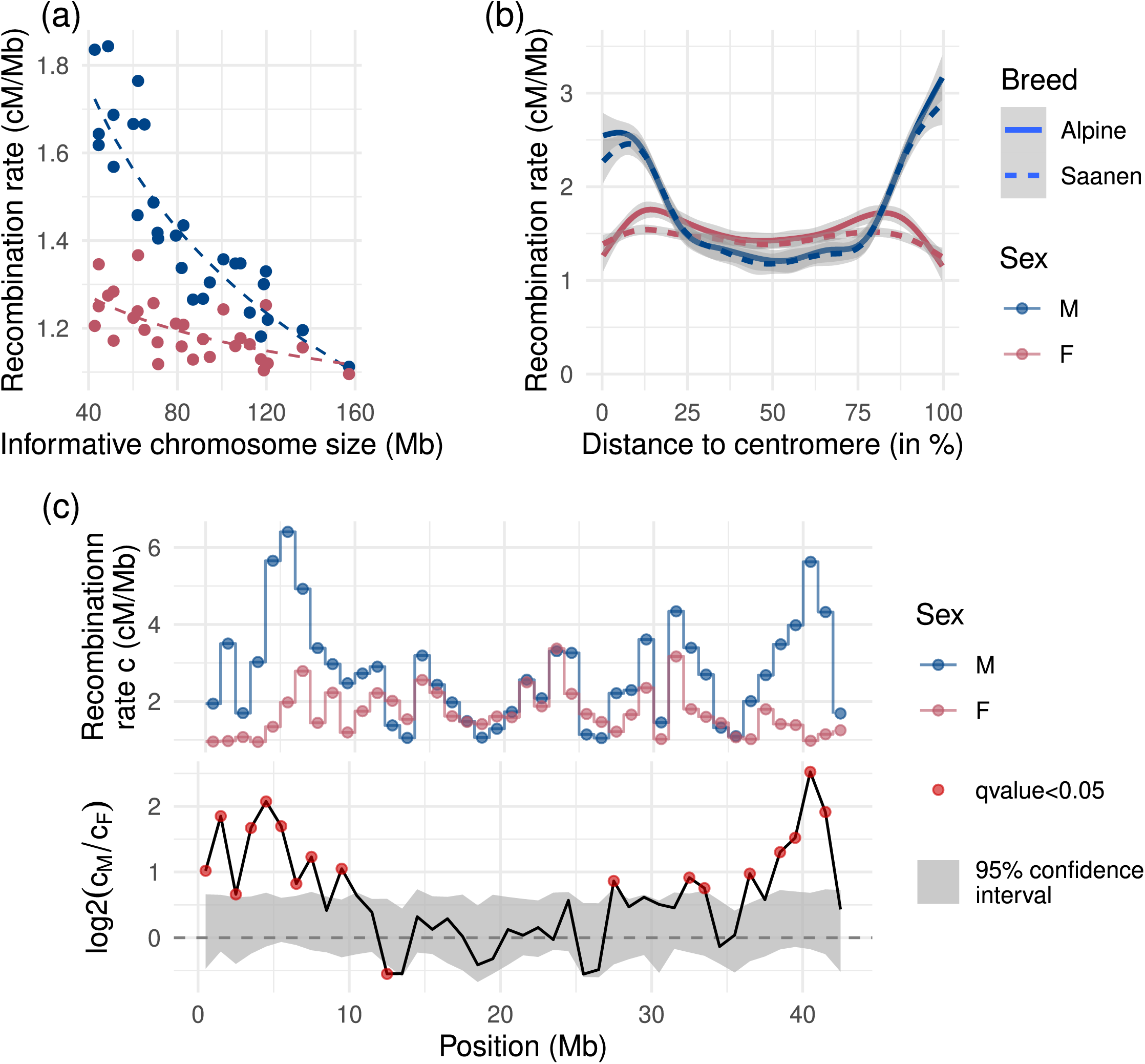
Estimated recombination rates for males and females at several scales. (a): Chromosomic recombination rate depending on chromosome size. (b): Bayesian estimates of the chromosome landscape of recombination rate depending on distance to centromere. All 29 chromosomes are acrocentric. Confidence intervals at 99%. (c): Recombination map on chromosome 25 at 1Mb resolution. Confidence interval under *H*_0_ estimated using sex permutations.

Chromosome landscape was calculated from 1Mb resolution Bayesian estimates of recombination rate: all positions in Mb were converted to positions relative to the centromere, and recombination rates of all chromosomes were studied depending on their position to the centromere. Considering that all goat autosomes are acrocentric, the beginning of all chromosomes (or centromeric tip) is located at 0%, and the end (or telomeric tip) is located at 100%. The results are presented in **Fig. 2b**. Females have a much flatter recombination pattern along the chromosomes compared to males, and both sexes seem to recombine as much in the middle of chromosomes.

### Local recombination rates

Recombination rates were estimated locally on 1Mb intervals on all autosomes (**See Additional File 4, Table S4**). Differences between sexes were estimated as the log ratio of male to female recombination rate and significance was assessed using its empirical null distribution calculated from permutations (see Methods). Recombination maps for each chromosome are available in **Additional File 5** (**Figures S18 to S46**), and **Fig. 2c** illustrates this inference on chromosome 25. It exhibits the general trend of the sex differences in recombination patterns where males recombine significantly more than females at the tips of the chromosomes. As all goat autosomes are acrocentric, this means that male recombination rate is higher not only at the telomeric end, but also at the centromeric end. A total of 871 intervals of the genome are significant for sex differences (out of 2477 intervals). Of the significant intervals, 559 (64% of the significant intervals) show higher recombination rate in males and are localized for the most part at both ends of chromosomes. The 312 remaining significant intervals (36% of the significant intervals) show higher recombination rate in females and are located in the middle of chromosomes. For a list of all significant intervals, see **Additional File 6, Table S5**.

### Genetic parameters and architecture of recombination phenotypes

#### Phenotype distributions

Two phenotypes were used to characterize the genetic basis of recombination in the goat: the individual Genome-wide Recombination Rate (GRR), a measure of recombination *intensity* and intra-chromosomal shuffling (*r̄*_*intra*_) a measure of the *distribution* of crossovers on chromosomes. This is because crossovers located in the middle of a chromosome lead to create more novel allelic combinations and therefore increase intra-chromosomal shuffling while crossovers located at the tips of a chromosome decrease intra-chromosomal shuffling.

GRR is defined as the per meiosis crossover count divided by the length of informative genome. The average value of GRR over all meioses is 1.39 cM per Mb. The distributions of GRR for different breeds and sexes are shown in **Figure S13** (**See Additional File 1**).

Intra-chromosomal shuffling *r̄*_*intra*_ was defined as stated in Material & Methods. The average value of *r̄*_*intra*_ is 0.011. The distributions of *r̄*_*intra*_ between breeds and sexes are shown in **Figure S14** (**See Additional File 1**). A linear model was fitted to estimate differences between populations. A representation of the estimates can be found in **Figure S15** (**See Additional File 1**). No significant difference was found between sexes (p-value = 0.12) but Alpine goats had significantly higher *r̄*_*intra*_ (p-value = 0.03). Furthermore, the interaction between breed and sex was found significant (female Alpine +4.01 × 10^−4^; p-value = 0.03), meaning that in average, Alpine females shuffled more than Alpine males, but Saanen females shuffled less than Saanen males.

#### Heritability estimates

Heritabilities of GRR and *r̄*_*intra*_ were estimated using an animal model taking into account repeated measures, only for the male phenotype. GRR heritability was estimated at 0.12±0.03 and was consistent in both breeds. *r̄*_*intra*_ heritability was estimated at 0.034±0.015 when both breeds were analysed together. No significant permanent environment effect was detected for any phenotype (**Table 2**).

**Table 2:**
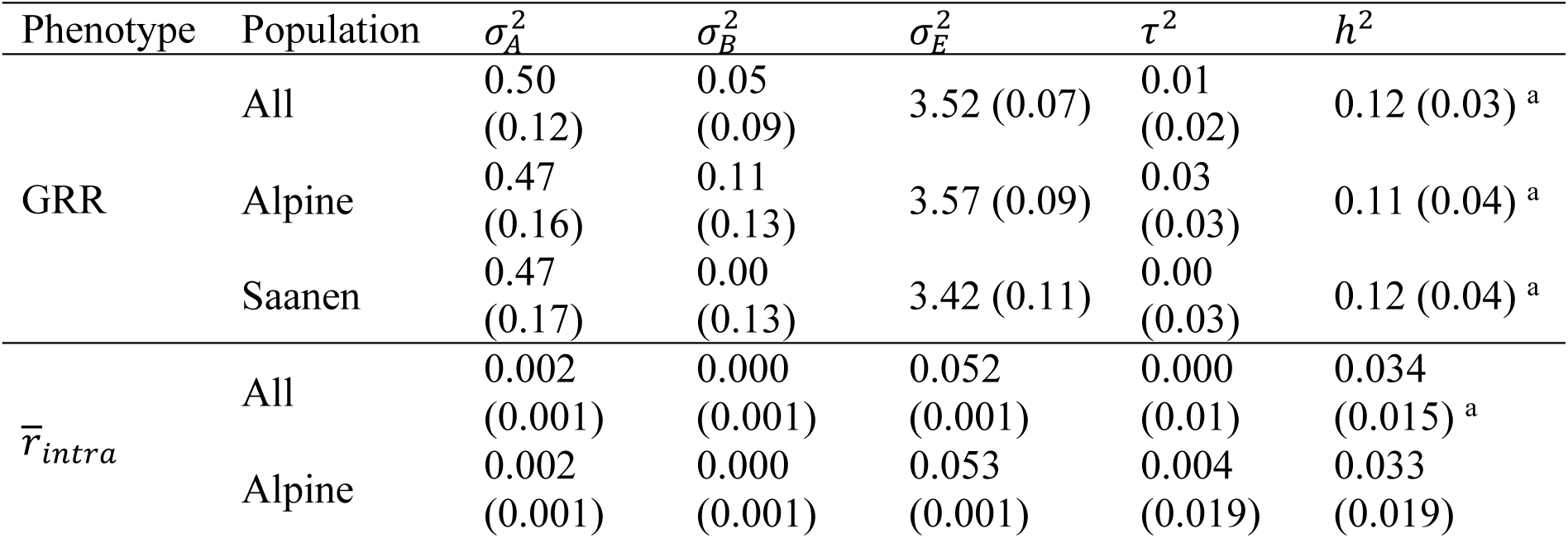
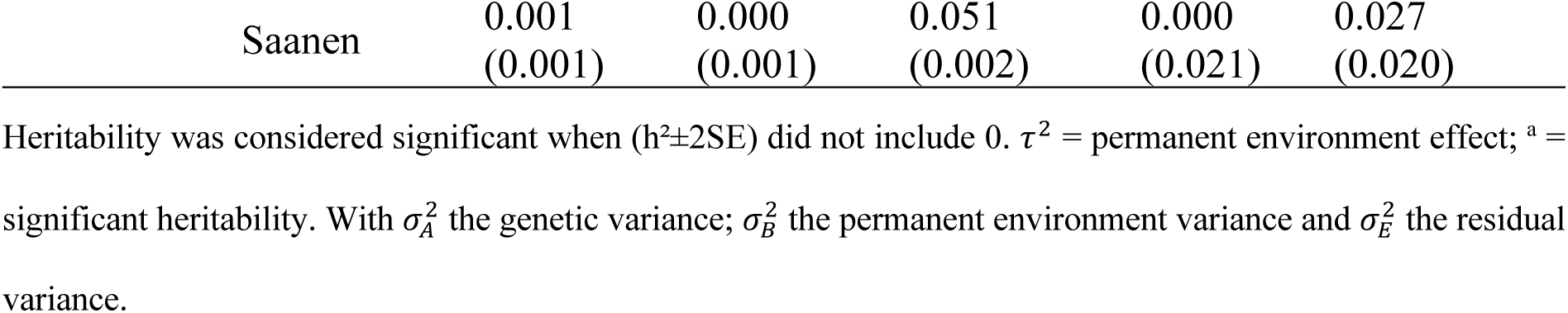
Variance components and heritability (SE) estimates for male genome-wide recombination rate and intra-chromosomal shuffling in a population of 843 Alpine and Saanen goats.

#### Genome-Wide Associations

The EBVs predicted from the animal model were used as phenotypes for the GWAS on GRR and *r̄*_*intra*_. GWAS results using imputed genotypes to the sequence level are illustrated in **Fig. 3** for Alpine and Saanen, on both phenotypes.

**Figure 3:**
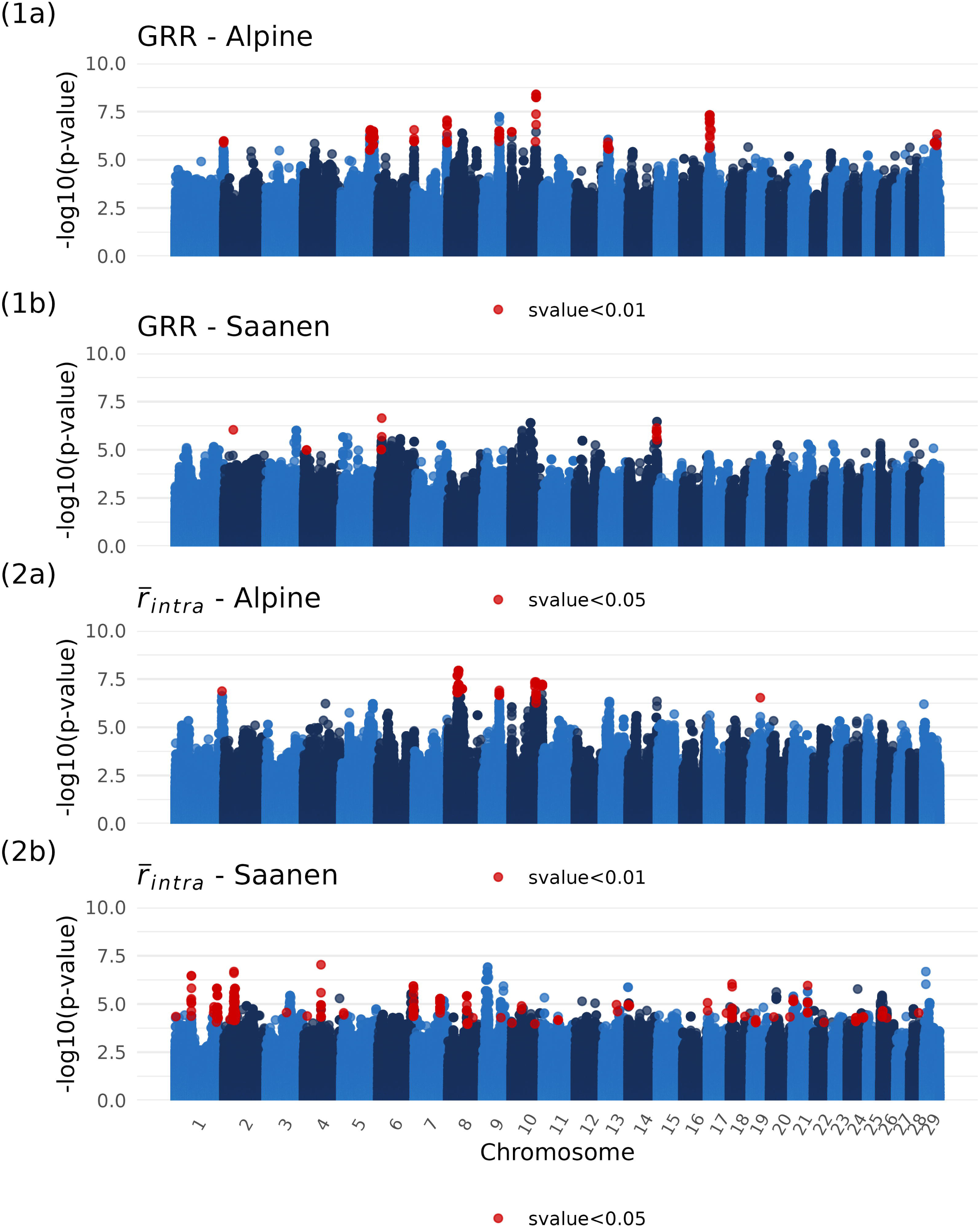
Manhattan plots of genome-wide association studies (GWAS) for male recombination phenotypes after imputation to sequence level. GRR and *r̄*_*intra*_ stand respectively for EBVs on male GRR and EBVs on male *r̄*_*intra*_.

The analyses on GRR revealed 15 significant QTLs in Alpine and 4 suggestive QTLs in Saanen. For Alpine, there were one QTL on chromosomes 1, 6, 7, 13 and 29; three QTLs on chromosomes 5 and 10; as well as two QTLs on chromosomes 9 and 17. For Saanen, there were one suggestive QTL on chromosomes 2, 4, 6 and 14. There is no overlap between the significant QTLs in Alpine and the suggestive QTLs in Saanen. Correlation of SNP effects between the two breeds for this phenotype was calculated using standard errors as weights. It amounted to 0.25.

The analyses on *r̄*_*intra*_ revealed 7 significant QTLs in Alpine and 41 suggestive QTLs in Saanen. Correlation of *β̂* between the two breeds is 0.04. For Alpine, there were one QTL on chromosomes 1, 9 and 19; as well as two QTLs on chromosomes 8 and 10. The lead variant on chromosome 9 is 5kb away from the closest significant QTL for GRR GWAS, and the one on chromosome 10 is 268bp away from its respective significant QTL for GRR. Correlation of the *β̂* between the GWAS for GRR and *r̄*_*intra*_ is 0.89. For Saanen, there were five QTLs on chromosome 1; two QTLs on chromosomes 2, 4, 6, 8, 14, 17, 18, 19, 20, 21 and 24; one QTL on chromosomes 3, 5, 7, 9, 11, 13, 22 and 28; as well as three QTLs on chromosome 10 and 26. Despite some QTLs being on the same chromosomes as for GRR, no suggestive region is within 2Mb of suggestive QTLs for GRR. Correlation between the two GWAS in Saanen is 0.92.

Considering the GWAS on GRR, the most significant QTL in Alpine is located on chromosome 17 at around 7Mb (lead variant s-value=0.00236) and is illustrated in **Fig. 4a**. In this region, 58 candidate genes can be found, including *SSH1* which scores 5/5. The significant SNPs are located directly within the *SSH1* gene, in an intronic region. The lead variant has a MAF of 0.49 in the population. The QTL illustrated in **Fig. 4b** contains significant SNPs in high LD in a region of 300kb and is rich in genes. It is the second most significant QTL (lead variant s-value=0.00238). Our standardised method for ranking candidate genes identifies 86 genes with a score of at least 1/5. The highest scoring (3/5) is *CARMIL3* because of its physical proximity to the lead variant. Among the candidates for this region are two genes previously associated with recombination: *RNF212B* (rank 17, score 1/5) and *REC8* (rank 51, score 1/5). The QTL illustrated in **Fig. 4c** is located at the end of chromosome 6 and is the 6^th^ most significant of the genome. It contains 36 candidate genes including *SPON2* (rank 1, score 4/5) and *RNF212* (rank 2, score 3/5).

**Figure 4:**
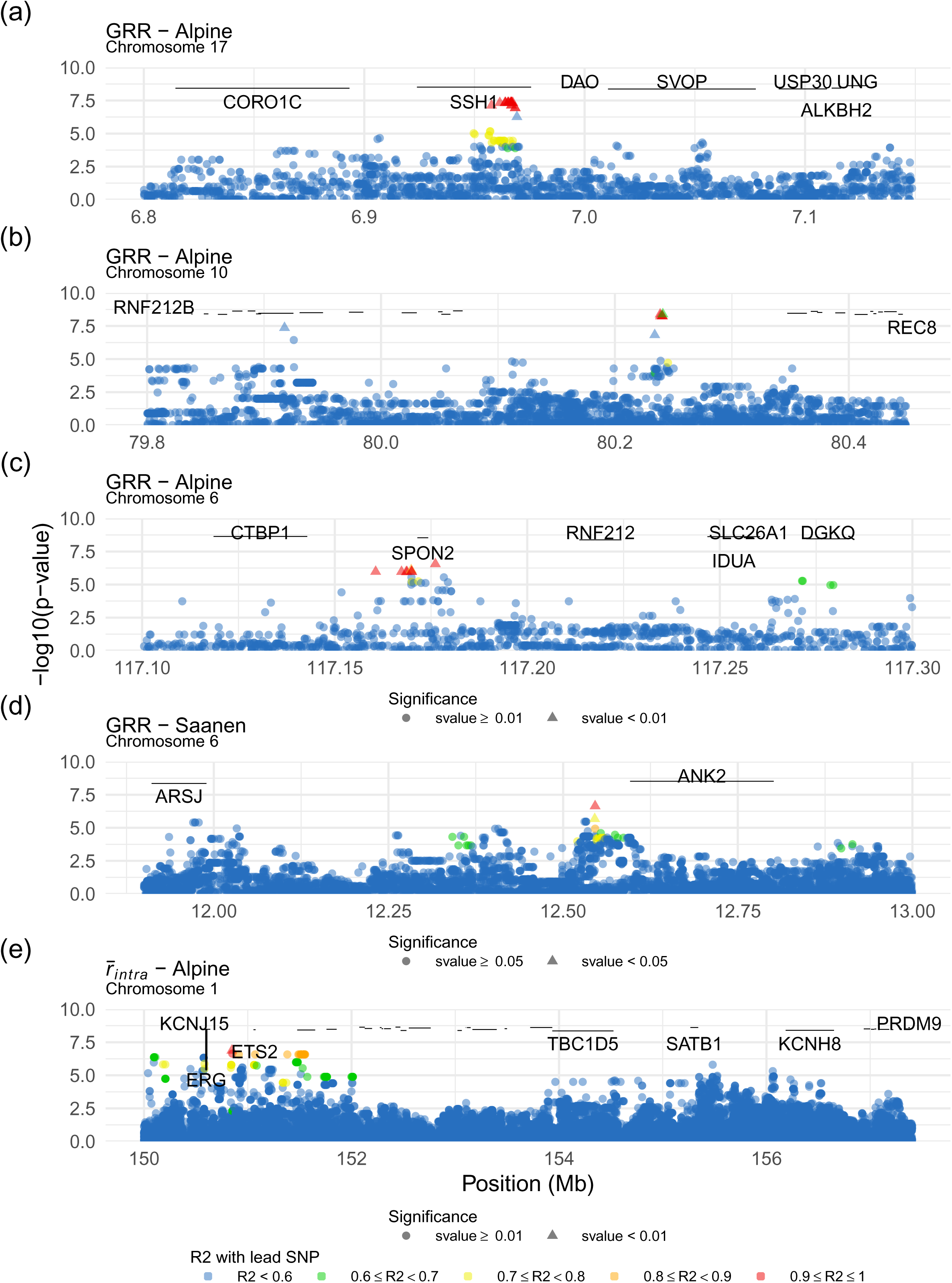
Zoom on GWAS Manhattan plots for regions of interest in both populations and phenotypes.

In Saanen, the most suggestive QTL can be found on chromosome 6 at around 12.5Mb (lead variant s-value=0.0326) and is illustrated in **Fig. 4d**. Several significant SNPs of the area fall within the *ANK2* gene which scores at 4/5 as a result. Eight other candidate genes can be found in the region. Other notable genes in both breeds can be found in Table 3. For exhaustive results for GWAS candidate genes in both breeds, see **Additional File 7**, **Tables 6 & 7**.

**Table 3:** Genes scoring above 4/5 for the GWAS on EBVs of male GRR in Alpine or Saanen.

| Breed | QTL of interest<br>(chromosome: position of<br>the lead variant) | Candidate gene (score/5) |
| --- | --- | --- |
| Alpine | 1:156523831 | KCNH8 (5/5) |
|  | 5:94902996 | GRIN2B (5/5) |
|  | 5:106271734 | B4GALNT3 (5/5) |
|  | 6:117176129 | SPON2 (4/5) |
|  | 7:105398227 | SNAP47 (4/5) |
|  | 9:53226533 | THEMIS (5/5) |
|  | 10:1686345 | EFCAB11 (5/5) |
|  | 13:17395272 | CCNY (4/5) |
|  | 17:6965601 | SSH1 (5/5) |
|  | 17:7730110 | GLTP (5/5) |
|  | 29:42944888 | FERMT3 (5/5) |
| Saanen | 2:29863833 | TNS1 (5/5) |
|  | 6:12545797 | ANK2 (4/5) |
|  | 14:91668757 | B3GAT2 (5/5) |

Considering the GWAS on *r̄*_*intra*_, the most significant QTL is the first signal of chromosome 8 in Alpine (lead variant s-value=0.0015). The gene *PTPRD*, which scores 4/5 for this region, has no known link with the recombination process. Another significant QTL can be found for this phenotype and breed at the end of chromosome 1 (**Fig. 4e**). The highest scoring candidate genes are *ERG* (4/5), *ETS2* (3/5) and *KCNJ15* (2/5), but none of them seem to be involved in recombination either. The region is 6Mb away from *PRDM9*. The most significant QTL for *r̄*_*intra*_ in Saanen can be found on chromosome 4 (lead variant s-value=0.0026). The gene *AVL9* scores 5/5 for this region but does not seem to be involved in the recombination process. For exhaustive candidate genes results for GWAS on *r̄*_*intra*_, see **Additional File 7, Tables 8 & 9**.

## Discussion

This study is a detailed analysis of the recombination process and its genetic architecture in a new species: the goat (*Capra Hircus*). It provides precise estimates of recombination rates along the genome, using a new statistical inference approach that accounts for different levels of uncertainty in crossover data. This analysis revealed that the genetic length of the ARS1 goat genome assembly is 35.1M (Alpine males) corresponding to a genome-wide recombination rate of 1.42cM/Mb. However, recombination was found to be highly variable along the genome, with rates varying from 0.21 to 10.32cM/Mb. An important feature of the recombination landscape is the large heterochiasmy in some genomic regions, mostly located at chromosome ends. This is consistent with patterns of recombination in a closely related species, the sheep *Ovis aries* [12,19].

In a second part of this study, we characterized the genetic architecture of recombination intensity and localisation through the study of two recombination phenotypes, GRR and *r̄*_*intra*_. This analysis revealed low but significant genetic components to these traits and identified several QTLs for both traits, some of which match loci found in other species while others are new. Overall, our study adds to the growing knowledge on recombination and will contribute to understanding the evolution of recombination, a fundamental trait for individual fitness and adaptive potential of populations. We now discuss the importance of our findings and the future research questions it suggests.

### Studying recombination from livestock breeding programmes data

Livestock breeding populations are good models to study the genetic architecture of the recombination process as they offer large datasets of genotyped individuals related through well characterised pedigrees. An important caveat of these datasets however lies in the unbalanced genotyping between sexes. In ruminant programs of genomic selection such as the one used here, male selection candidates are typically all genotyped while females are much less likely to be, although this tends to change with the decrease in genotyping costs. In our dataset, the number of dams was very small. We still managed to correctly characterise heterochiasmy along the genome, but the number of dams was too small to study the genetic architecture of recombination in this sex. We expect the number of genotyped females to grow in the future which will allow studying genetic determinants of heterochiasmy in this species. However, it should be noted that offspring size will still typically be smaller for females, so that taking into account uncertainty as we did here will remain relevant.

### Recombination landscape of the goat genome

Our study employs a similar strategy as previous studies in animal populations [8,10–12,16,18–20,22,46], but we aimed at better accounting for the uncertainty in crossover data. The first source of uncertainty arises from the unprecise localisation of crossovers due to the genotyping density, the second by the homozygosity of individuals in the pedigree, and the third concerns only female recombination rate estimation, as our available female data is much scarcer. To tackle the first problem, [12] proposed a random sampling approach of crossover locations to average recombination rate estimation over crossover uncertainty. Here, we used the same strategy but extended it further to better identify genomic regions with significant differences in recombination rates between sexes. To tackle the second problem, we made use of an information provided by the software used to phase individuals (Yapp) that explicitly identifies regions of the genome where meioses are uninformative for crossover detection. In order to integrate both sources of uncertainty, we leveraged the Bayesian Gamma-Poisson model of [12] that allows to incorporate this information naturally. One caveat of this model is that it is much harder to implement if one wants to add a hierarchical level to account for differences between groups (*e.g.* sex) but with non-null covariances. Recent developments around Poisson-Log-Normal models [47] could be used for this purpose (see e.g. [48] in the context of recombination rate inference) but this will require further developments, in particular to better account for the longitudinal correlation of recombination rates along the genome. Finally, we proposed a permutation method to assess the significance of heterochiasmy along chromosomes. This method allows to draw conclusions from a lowly informative dataset in terms of sex differences, which could not have been confidently untangled otherwise. We believe it could be useful in future studies on heterochiasmy.

The average recombination rate of the goat genome (1.42cM/Mb varying from 1.12 to 1.71cM/Mb among chromosomes) is very similar to other ruminants (e.g. 1.5cM/Mb in sheep [12], 0.98cM/Mb in cattle [46]). A striking feature of its landscape is a strong heterochiasmy that we could reveal even if the number of female meioses available in our datasets was relatively low. This heterochiasmy is mostly concentrated in a few megabases at the chromosome ends, irrespective of the total chromosome size. As a consequence, the difference in average recombination rate between sexes is smaller for large chromosomes than small ones (**Fig. 2a**). The biological causes underlying this heterochiasmy are still unknown. Further research combining genetics studies with cytogenetics and functional studies will most likely be required to unravel them.

Patterns of recombination and heterochiasmy in the goat match those found in sheep [12,19,48] and cattle [8,10,11] although maybe to a lesser extent for this last species. Given that these three species have highly colinear genomes, it seems possible to make a quantitative comparison of recombination patterns along ruminant chromosomes to reveal how recombination evolved with speciation.

### Genetic parameters and architecture of recombination phenotypes

We studied two different phenotypes: the Genome-Wide Recombination Rate (GRR) which is a measure of recombination intensity, and intra-chromosomal shuffling (*r̄*_*intra*_) which depends on recombination intensity too but also measures differences in the localisation of crossovers. As expected, we observed a high correlation between the two phenotypes (0.62). However, despite strong correlation of the GWAS results on GRR and *r̄*_*intra*_, their genetic architectures were found only partially overlapping, illustrating that they are probably capturing different underlying biological processes (for example those involved in Double Strand Break (DSB) localisation from those involved in DSB repair).

Heritability for male GRR was estimated at 0.12 with similar results for each breed, and similar to those found in other ruminant species for this phenotype (ACC or GRR): *e.g.* 0.22 for Dutch Holstein-Friesan males in cattle [6], 0.13 in several cattle breeds [9], 0.12 for Soay sheep [19] and 0.23 for Lacaune sheep [12].

The heritability for male *r̄*_*intra*_ was estimated at 0.034 and consistent between the two breeds, although only slightly significantly different from zero (SE=0.015). Heritability for this phenotype seems to be smaller than in other species as [18] estimated it at 0.11 in salmon, a species with *PRDM9* hotspots, and [22] estimated it at 0.14 in sparrows, a species without *PRDM9* hotspots.

The animal models used for estimating genetic parameters also allow to predict the additive genetic values (EBVs) of every animal in the pedigree. These EBVs on male GRR and *r̄*_*intra*_ were used as phenotypes for our GWAS. This permits to have only one measure of phenotype for each individual, and therefore to run analyses using the fast computing procedures required for whole genome sequence imputed datasets. Using EBVs as phenotypes for GWAS is sometimes discouraged for different reasons. First, to avoid too much shrinkage in the phenotypes: ideally all animals should have either own and/or direct offspring phenotypes. Here, out of the 843 individuals in the GWAS, only 323 (38%) did not have any own phenotypes. These were all females, as they obviously cannot be directly measured for male recombination phenotypes. Among these females, only 43 (5% of all animals) were not directly related to a male with own information for these phenotypes, *i.e.* had a coefficient of relationship smaller than ½ with measured samples. So overall, our dataset meets this first requirement. Another caveat of using EBVs as phenotype is that their precisions can sometimes vary widely between individuals, especially when there are large differences in their number of repeated measures. In our case, the posterior variances of EBVs ranged from 2.6 to 7.1, *i.e.* a factor smaller than 3 between the most and least precise predictions. Also, although precisions of females EBVs tended to be smaller than males their distribution was unimodal (**See Additional File 1, Figures S16 & S17**). Given this homogeneity in the precision of EBVs, we did not deem deregression, a procedure that comes with its own caveats, necessary. Overall, for this dataset we believe that phenotypic records are sufficiently informative and homogeneously distributed among animals to use EBVs as phenotypes. Finally, we note that we used the pedigree relationship matrix to derive EBVs while the GRM was used in the GWAS, in addition to using the LOCO methodology, avoiding potential convergence issues (see **Methods**).

GWAS results for male GRR and *r̄*_*intra*_ showed low genetic correlation between Alpine and Saanen. Despite identical heritability results, no QTL were found in common between the two breeds. Besides, the correlation of the *β̂* for the SNPs that are significant in at least one breed is only 0.25 for GRR and 0.04 for *r̄*_*intra*_. This suggests strong differences in the underlying genetic determinants of recombination between these two breeds, although they are both European breeds originating from the Alps and closely related in a worldwide diversity panel [49]. Despite this relatively close genetic proximity, those breeds have been shown to have different genetic architecture for other traits. For example, [50] found genetic heterogeneity in the determinism of dairy-related traits between the two breeds, despite similar breeding goals. Our results add to the literature that suggests that genome-wide recombination landscapes tend to be mostly shared between populations while the underlying genetic determinants of the process can be quite different, *e.g.* [12] in sheep, [15,16] in pigs, although results in cattle tend to show they are more conserved [10].

Candidate genes associated with QTL loci in the GWAS were prioritised by implementing a scoring methodology using criteria based on physical and genetic distance to significant SNPs. This methodology was developed in order to provide an objective measure for the investigation of candidate genes underlying GWAS peaks and quickly highlight strong candidates in regions rich in annotations. For instance, 13Mb on chromosome 5 contain three different peaks in rapid succession and 141 genes are annotated in the region. The scoring allowed to quickly identify the most promising genes and order them all depending on their physical and genetic proximity to each peak. Genes involved in meiosis in other species were searched for in a second step to identify functional candidates at QTLs among the positional candidates.

The GWAS results for GRR in Alpine goats show a signal towards the end of chromosome 6, in very close proximity to the *RNF212* gene. *RNF212* has been found to be associated with GRR in humans [51], cattle [6,9] and sheep [12,19]. It has also been proved that *RNF212* is involved in and is necessary for meiosis in mammals by playing a central role to the designation of crossover sites and the formation of crossover-specific recombination complexes [52,53]. Yet it only scores second candidate for the signal on chromosome 6 because it does not directly contain the significant SNPs. Of course, this is still consistent with a regulatory variant affecting *RNF212* expression.

One signal with significant SNPs as far as 330kb apart can be found at the end of chromosome 10 between 79.9 and 80.3Mb in the Alpine breed (**Fig. 4b**). The genes *RNF212B* and *REC8* are in the region. They were previously suspected to be responsible for signals in sheep [12,19] and cattle [6,9] but the studies had difficulty attributing the signal to one of the two genes due to a lack of resolution. Among our candidates, *RNF212B* ranks 17^th^ and *REC8* ranks 51^st^, as the region is very rich in annotated genes. Functionally, both these genes make very good candidates. On the one hand, *RNF212B* is a homolog of *RNF212*, this gene’s involvement in recombination being described just above, and the two proteins *rnf212* and *rnf212b* have been showed to interact with each other and to be necessary for their stability during the recombination process [54]. On the other hand, REC8 is also known to be involved in meiotic recombination [55] through DSB formation [56] as well as in implementing homologue bias during DSB repairs [57–59]. But the gene richness of the region (62 genes in 690kb), their relative physical distance to the lead variant and the fact that, surprisingly, the main part of the signal is located in an area of 250kb completely devoid of annotated protein coding genes, makes discriminating the two candidates, or even determining if they are good candidates, quite difficult.

Among significant QTLs, the most significant signal was found in Alpine on chromosome 17 around 7Mb (**Fig. 4a**). The signal falls directly within the gene *SSH1*, which scored 5/5 according to our ranking method. This gene belongs to the slingshot phosphatase family, constituted of three genes [60]. Investigation of the GTEx database [61,62] showed that it is expressed in testis and automatic annotations [63,64] indicate it may be involved in chromatin remodelling mechanisms. Furthermore, its paralog *SSH2* showed protein-enrichment in a study aiming to predict mouse meiosis-essential genes [65] and is known to be involved in later spermatogenesis [66]. But current knowledge of these genes’ functions as well as the strong signal found for a QTL within *SSH1* suggest this family may be involved in recombination.

When finding QTLs of interest at the end of chromosome 1 in Alpine, both for GRR and *r̄*_*intra*_, one could have assumed that these signals were associated with *PRDM9*, known regulator of recombination hotspots [67,68] and located at the very end of chromosome 1 in goats (**See Additional File 3**, **Table 1**). Therefore, it was surprising to see that *PRDM9* actually scores very low for the GRR signal (1/5 score) due to its physical distance from the significant QTL and is even too far from the *r̄*_*intra*_ signal to gain any score point. It could be explained that no association was found between *PRDM9* and GRR because GRR is a phenotype of intensity of recombination, and *PRDM9* regulates the location of recombination. But *r̄*_*intra*_ is a recombination phenotype linked with the position of crossovers and the association was even more expected, especially considering that [17] found a clear signal for *PRDM9* in their own GWAS on pig *r̄*_*intra*_ phenotype. Furthermore, the third highest ranking candidate gene in goats for *r̄*_*intra*_ is *KCNJ15* (2/5 score and **Fig. 4e**). Although this gene seems to have no link with recombination, an association for this gene with GRR has previously been found in sheep [12]. Furthermore, [6] found signal for the genes *KCNJ2* and *KCNJ16*, paralogs of *KCNJ15*, in their GWAS for GRR. Thus, further investigation on the involvement of *KCNJ15* in recombination may be relevant.

## Conclusions

In this study, we have found that recombination patterns are very alike between the Alpine and Saanen breeds, but heterochiasmy is strongly present, manifested at the chromosome ends. The recombination pattern we observed is well conserved with closely related species like sheep and cattle. We also showed that male GRR and male *r̄*_*intra*_ are heritable and that this value is conserved in the two breeds. Heritability for GRR is concordant with results found in sheep and cattle, but goat heritability for *r̄*_*intra*_ seems to be lower than in the other species in which it was estimated. GWAS results showed low correlation between the two breeds, for both phenotypes, and it seems that despite similar recombination patterns and same heritability for recombination traits, the underlying genetic architecture of recombination is different between Alpine and Saanen. Candidate genes found in GWAS are partially in common with what was previously found in literature, with some novel genes. This supports the hypothesis that genetic control for recombination is well conserved among many lowly-related species, but also that the genes involved in this control can evolve quickly.

## Declarations

### Ethics approval and consent to participate

Not applicable

### Consent for publication

Not applicable

### Availability of data and materials

Initial genotypes and pedigree were provided by Capgenes and INRAE. Their access is restricted by the provider. Crossover data and informativity data, derived from these initial datasets, as well as genotypes of the phenotyped parents and scripts for analysis are available at https://doi.org/10.57745/HPYLLT. Sequence data for the imputation reference panel can be obtained from the VarGoats consortium: https://www.goatgenome.org/vargoats.html.

## Authors’ contributions

All authors participated in conceptualizing this study and to the data curation and analyses. All authors developed methodology for this study and administered the project. RR provided the initial dataset as resources for the project. BS and AE did programming and software development to produce results. BS and RR supervised the project. All authors validated the results. AE and BS did visualisation work through diverse presentations and wrote the original draft. All authors reviewed, edited and approved the final manuscript.

## Acknowledgements

Many numerical analyses were performed on the genobioinfo bioinformatics platform Toulouse Midi-Pyrénées (Bioinfo Genotoul). We are thankful to Capgenes and the INRAE P3R unit (https://doi.org/10.15454/1.5483259352597417E12) for providing access to SNP array genotypes, and to the VarGoats consortium for early access to whole genome sequencing data. We would like to thank Susan Johnston, Tristan Mary-Huard and Leopoldo Sanchez Rodriguez for fruitful discussions during the production of these results.

## Additional Files

### Additional File 1 Figure S1

File format: PDF

Title: Position of (b)ovine (top) and goat (bottom) chip SNPs on both the RH map and the ARS1 assembly, on chromosomes 2 and 18.

Description: Goat SNPs have been filtered using the longest increasing subsequence algorithm, and only remaining SNPs have been represented.

### Additional File 1 Figure S2

File format: PDF

Title: Crossover distribution on the 35-45 Mb area of chromosome 1.

Description: Red lines = position of conserved synteny breaks.

### Additional File 1 Figure S3

File format: PDF

Title: Additive SNP and sample call-rate cumulative distributions.

Description: Left: SNP call-rate; right: sample call-rate. Black line: chosen threshold=0.01.

### Additional File 1 Figure S4

File format: PDF

Title: Distribution of the distance between two crossovers per chromosome per meiosis. Description: Red line: chosen threshold=5Mb

### Additional File 1 Figure S5

File format: PDF

Title: Density distribution of the Maximum Likelihood estimates of recombination rates on 1Mb intervals of the genome.

Description: Black: gamma law; extracted parameters α=3.08 & β=2.20.

### Additional File 1 Figure S6

File format: PDF

Title: Distribution of crossover detection interval sizes.

### Additional File 1 Figure S7

File format: PDF

Title: Distribution of all permuted recombination rate log ratio for all genomic intervals.

### Additional File 1 Figure S8

File format: PDF

Title: SNP distribution of sequencing and variant calling quality parameters.

Description: A: DP parameter; B: FS parameter; C: VQSLOD parameter.

### Additional File 1 Figure S9

File format: PDF

Title: R² evolution according to allele frequency.

Description: (A) Before genotype imputation. Mean R²=0.928. (B) After genotype imputation. Mean R²=0.990.

### Additional File 1 Figure S10

File format: PDF

Title: Genotype concordance according to allele frequency.

Description: (A) Before genotype imputation. Mean genotype concordance=0.974. (B) After genotype imputation. Mean genotype concordance=0.997.

### Additional File 1 Figure S11

File format: PDF

Title: Distribution the proportion of the autosomal genome informative for crossovers in parents (M=males vs F=females).

### Additional File 1 Figure S12

File format: PDF

Title: Comparison of recombination map size estimates between populations.

Description: Confidence interval = *μ*±2×*standard error*.

### Additional File 1 Figure S13

File format: PDF

Title: Distribution of genome-wide recombination rates (GRR) in meioses.

### Additional File 1 Figure S14

File format: PDF

Title: Distribution of intra-chromosomal shuffling in meioses.

### Additional File 1 Figure S15

File format: PDF

Title: Comparison of intra-chromosomal shuffling estimates between populations. Description: Confidence interval = *μ*±2×*standard error*.

### Additional File 1 Figure S16

### Additional File 1 Figure S17

### Additional File 2 Table S1

### File format: XLS

Title: SNPs within synteny breakpoints

Description: File listing all 123 SNPs removed from dataset after identifying synteny breakpoints with human genome on ARS1 assembly.

### Additional File 3 Table S2

File format: XLS

Title: BLAST results for the bovine gene sequence RNF212 on ARS1 assembly.

### Additional File 3 Table S3

File format: XLS

Title: BLAST results for the bovine gene sequence PRDM9 on ARS1 assembly.

### Additional File 4 Table S4

File format: XLS

Title: Sex-specific estimated recombination rates for each 1Mb interval of the autosomal goat genome.

### Additional File 5 Figure S18

File format: PDF

Title: Estimated recombination rates for males and females on chromosome 1, at 1Mb resolution.

Description: Log ratio between male and female recombination rates were plotted in the bottom panel. Confidence interval of log ratio under null hypothesis appears in grey. Intervals found significant for sex differences are marked with a red dot on the bottom panel.

### Additional File 5 Figure S19

File format: PDF

Title: Estimated recombination rates for males and females on chromosome 2, at 1Mb resolution.

Description: Log ratio between male and female recombination rates were plotted in the bottom panel. Confidence interval of log ratio under null hypothesis appears in grey. Intervals found significant for sex differences are marked with a red dot on the bottom panel.

### Additional File 5 Figure S20

File format: PDF

Title: Estimated recombination rates for males and females on chromosome 3, at 1Mb resolution.

Description: Log ratio between male and female recombination rates were plotted in the bottom panel. Confidence interval of log ratio under null hypothesis appears in grey. Intervals found significant for sex differences are marked with a red dot on the bottom panel.

### Additional File 5 Figure S21

File format: PDF

Title: Estimated recombination rates for males and females on chromosome 4, at 1Mb resolution.

Description: Log ratio between male and female recombination rates were plotted in the bottom panel. Confidence interval of log ratio under null hypothesis appears in grey. Intervals found significant for sex differences are marked with a red dot on the bottom panel.

### Additional File 5 Figure S22

File format: PDF

Title: Estimated recombination rates for males and females on chromosome 5, at 1Mb resolution.

Description: Log ratio between male and female recombination rates were plotted in the bottom panel. Confidence interval of log ratio under null hypothesis appears in grey. Intervals found significant for sex differences are marked with a red dot on the bottom panel.

### Additional File 5 Figure S23

File format: PDF

Title: Estimated recombination rates for males and females on chromosome 6, at 1Mb resolution.

Description: Log ratio between male and female recombination rates were plotted in the bottom panel. Confidence interval of log ratio under null hypothesis appears in grey. Intervals found significant for sex differences are marked with a red dot on the bottom panel.

### Additional File 5 Figure S24

File format: PDF

Title: Estimated recombination rates for males and females on chromosome 7, at 1Mb resolution.

Description: Log ratio between male and female recombination rates were plotted in the bottom panel. Confidence interval of log ratio under null hypothesis appears in grey. Intervals found significant for sex differences are marked with a red dot on the bottom panel.

### Additional File 5 Figure S25

File format: PDF

Title: Estimated recombination rates for males and females on chromosome 8, at 1Mb resolution.

Description: Log ratio between male and female recombination rates were plotted in the bottom panel. Confidence interval of log ratio under null hypothesis appears in grey. Intervals found significant for sex differences are marked with a red dot on the bottom panel.

### Additional File 5 Figure S26

File format: PDF

Title: Estimated recombination rates for males and females on chromosome 9, at 1Mb resolution.

Description: Log ratio between male and female recombination rates were plotted in the bottom panel. Confidence interval of log ratio under null hypothesis appears in grey. Intervals found significant for sex differences are marked with a red dot on the bottom panel.

### Additional File 5 Figure S27

File format: PDF

Title: Estimated recombination rates for males and females on chromosome 10, at 1Mb resolution.

Description: Log ratio between male and female recombination rates were plotted in the bottom panel. Confidence interval of log ratio under null hypothesis appears in grey. Intervals found significant for sex differences are marked with a red dot on the bottom panel.

### Additional File 5 Figure S28

File format: PDF

Title: Estimated recombination rates for males and females on chromosome 11, at 1Mb resolution.

Description: Log ratio between male and female recombination rates were plotted in the bottom panel. Confidence interval of log ratio under null hypothesis appears in grey. Intervals found significant for sex differences are marked with a red dot on the bottom panel.

### Additional File 5 Figure S29

File format: PDF

Title: Estimated recombination rates for males and females on chromosome 12, at 1Mb resolution.

Description: Log ratio between male and female recombination rates were plotted in the bottom panel. Confidence interval of log ratio under null hypothesis appears in grey. Intervals found significant for sex differences are marked with a red dot on the bottom panel.

### Additional File 5 Figure S30

File format: PDF

Title: Estimated recombination rates for males and females on chromosome 13, at 1Mb resolution.

Description: Log ratio between male and female recombination rates were plotted in the bottom panel. Confidence interval of log ratio under null hypothesis appears in grey. Intervals found significant for sex differences are marked with a red dot on the bottom panel.

### Additional File 5 Figure S31

File format: PDF

Title: Estimated recombination rates for males and females on chromosome 14, at 1Mb resolution.

Description: Log ratio between male and female recombination rates were plotted in the bottom panel. Confidence interval of log ratio under null hypothesis appears in grey. Intervals found significant for sex differences are marked with a red dot on the bottom panel.

### Additional File 5 Figure S32

File format: PDF

Title: Estimated recombination rates for males and females on chromosome 15, at 1Mb resolution.

Description: Log ratio between male and female recombination rates were plotted in the bottom panel. Confidence interval of log ratio under null hypothesis appears in grey. Intervals found significant for sex differences are marked with a red dot on the bottom panel.

### Additional File 5 Figure S33

Title: Estimated recombination rates for males and females on chromosome 16, at 1Mb resolution.

Description: Log ratio between male and female recombination rates were plotted in the bottom panel. Confidence interval of log ratio under null hypothesis appears in grey. Intervals found significant for sex differences are marked with a red dot on the bottom panel.

### Additional File 5 Figure S34

File format: PDF

Title: Estimated recombination rates for males and females on chromosome 17, at 1Mb resolution.

Description: Log ratio between male and female recombination rates were plotted in the bottom panel. Confidence interval of log ratio under null hypothesis appears in grey. Intervals found significant for sex differences are marked with a red dot on the bottom panel.

### Additional File 5 Figure S35

Title: Estimated recombination rates for males and females on chromosome 18, at 1Mb resolution.

Description: Log ratio between male and female recombination rates were plotted in the bottom panel. Confidence interval of log ratio under null hypothesis appears in grey. Intervals found significant for sex differences are marked with a red dot on the bottom panel.

### Additional File 5 Figure S36

File format: PDF

Title: Estimated recombination rates for males and females on chromosome 19, at 1Mb resolution.

Description: Log ratio between male and female recombination rates were plotted in the bottom panel. Confidence interval of log ratio under null hypothesis appears in grey. Intervals found significant for sex differences are marked with a red dot on the bottom panel.

### Additional File 5 Figure S37

Title: Estimated recombination rates for males and females on chromosome 20, at 1Mb resolution.

Description: Log ratio between male and female recombination rates were plotted in the bottom panel. Confidence interval of log ratio under null hypothesis appears in grey. Intervals found significant for sex differences are marked with a red dot on the bottom panel.

### Additional File 5 Figure S38

File format: PDF

Title: Estimated recombination rates for males and females on chromosome 21, at 1Mb resolution.

Description: Log ratio between male and female recombination rates were plotted in the bottom panel. Confidence interval of log ratio under null hypothesis appears in grey. Intervals found significant for sex differences are marked with a red dot on the bottom panel.

### Additional File 5 Figure S39

File format: PDF

Title: Estimated recombination rates for males and females on chromosome 22, at 1Mb resolution.

Description: Log ratio between male and female recombination rates were plotted in the bottom panel. Confidence interval of log ratio under null hypothesis appears in grey. Intervals found significant for sex differences are marked with a red dot on the bottom panel.

### Additional File 5 Figure S40

File format: PDF

Title: Estimated recombination rates for males and females on chromosome 23, at 1Mb resolution.

Description: Log ratio between male and female recombination rates were plotted in the bottom panel. Confidence interval of log ratio under null hypothesis appears in grey. Intervals found significant for sex differences are marked with a red dot on the bottom panel.

### Additional File 5 Figure S41

File format: PDF

Title: Estimated recombination rates for males and females on chromosome 24, at 1Mb resolution.

Description: Log ratio between male and female recombination rates were plotted in the bottom panel. Confidence interval of log ratio under null hypothesis appears in grey. Intervals found significant for sex differences are marked with a red dot on the bottom panel.

### Additional File 5 Figure S42

File format: PDF

Title: Estimated recombination rates for males and females on chromosome 25, at 1Mb resolution.

Description: Log ratio between male and female recombination rates were plotted in the bottom panel. Confidence interval of log ratio under null hypothesis appears in grey. Intervals found significant for sex differences are marked with a red dot on the bottom panel.

### Additional File 5 Figure S43

File format: PDF

Title: Estimated recombination rates for males and females on chromosome 26, at 1Mb resolution.

Description: Log ratio between male and female recombination rates were plotted in the bottom panel. Confidence interval of log ratio under null hypothesis appears in grey. Intervals found significant for sex differences are marked with a red dot on the bottom panel.

### Additional File 5 Figure S44

File format: PDF

Title: Estimated recombination rates for males and females on chromosome 27, at 1Mb resolution.

Description: Log ratio between male and female recombination rates were plotted in the bottom panel. Confidence interval of log ratio under null hypothesis appears in grey. Intervals found significant for sex differences are marked with a red dot on the bottom panel.

### Additional File 5 Figure S45

File format: PDF

Title: Estimated recombination rates for males and females on chromosome 28, at 1Mb resolution.

Description: Log ratio between male and female recombination rates were plotted in the bottom panel. Confidence interval of log ratio under null hypothesis appears in grey. Intervals found significant for sex differences are marked with a red dot on the bottom panel.

### Additional File 5 Figure S46

File format: PDF

Title: Estimated recombination rates for males and females on chromosome 29, at 1Mb resolution.

Description: Log ratio between male and female recombination rates were plotted in the bottom panel. Confidence interval of log ratio under null hypothesis appears in grey. Intervals found significant for sex differences are marked with a red dot on the bottom panel.

### Additional File 6 Table S5

File format: XLS

Title: Significant 1Mb intervals for sex differences in recombination rate.

### Additional File 7 Table S6

File format: PDF

Title: Candidate genes for GWAS on EBVs for male genome-wide recombination rate in Alpine goats.

Description: All genes scoring at least 1 as candidates are showed. For each QTL of interest, genes have been sorted from higher scoring to lower scoring. Manual annotation has been added on genes previously found in GWAS for recombination phenotypes in ruminants, referencing the corresponding papers.

### Additional File 7 Table S7

File format: PDF

Title: Candidate genes for GWAS on EBVs for male genome-wide recombination rate in Saanen goats.

Description: All genes scoring at least 1 as candidates are showed. For each QTL of interest, genes have been sorted from higher scoring to lower scoring. Manual annotation has been added on genes previously found in GWAS for recombination phenotypes in ruminants, referencing the corresponding papers.

### Additional File 7 Table S8

File format: PDF

Title: Candidate genes for GWAS on EBVs for male intra-chromosomal shuffling in Alpine goats.

Description: All genes scoring at least 1 as candidates are showed. For each QTL of interest, genes have been sorted from higher scoring to lower scoring. Manual annotation has been added on genes previously found in GWAS for recombination phenotypes in ruminants, referencing the corresponding papers.

### Additional File 7 Table S9

File format: PDF

Title: Candidate genes for GWAS on EBVs for male intra-chromosomal shuffling in Saanen goats.

Description: All genes scoring at least 1 as candidates are showed. For each QTL of interest, genes have been sorted from higher scoring to lower scoring. Manual annotation has been added on genes previously found in GWAS for recombination phenotypes in ruminants, referencing the corresponding papers.

