## Additional File 1 for "Genome landscape and genetic architecture of recombination in domestic goats (*Capra Hircus*)"

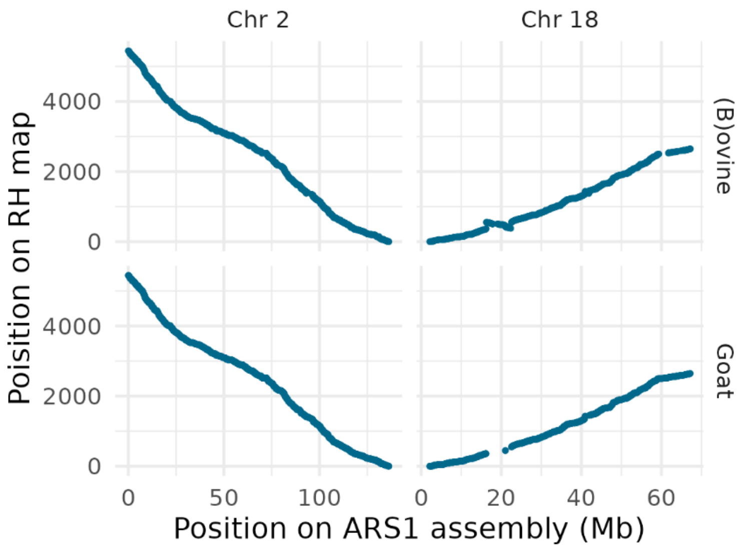


**Figure S1:** Position of (b)ovine (top) and goat (bottom) chip SNPs on both the RH map and the ARS1 assembly, on chromosomes 2 and 18. Goat SNPs have been filtered using the longest increasing subsequence algorithm, and only remaining SNPs have been represented.

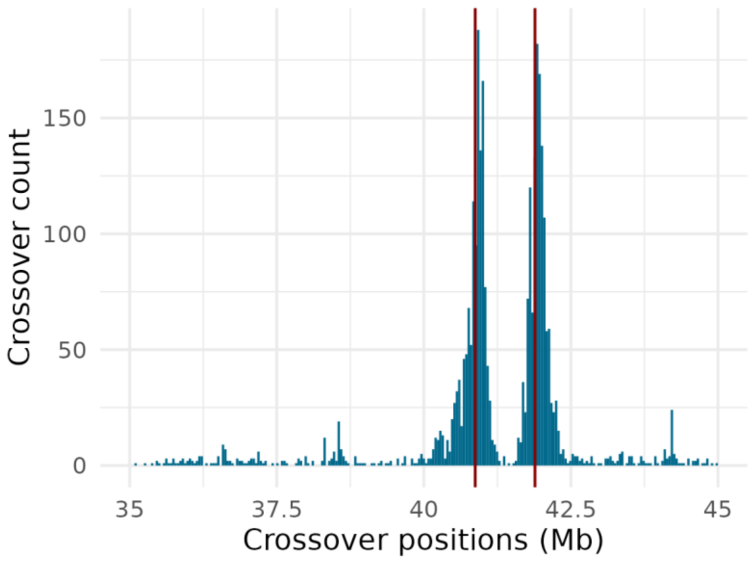


**Figure S2 :** Crossover distribution on the 35-45 Mb area of chromosome 1. Red lines = position of conserved synteny breaks.

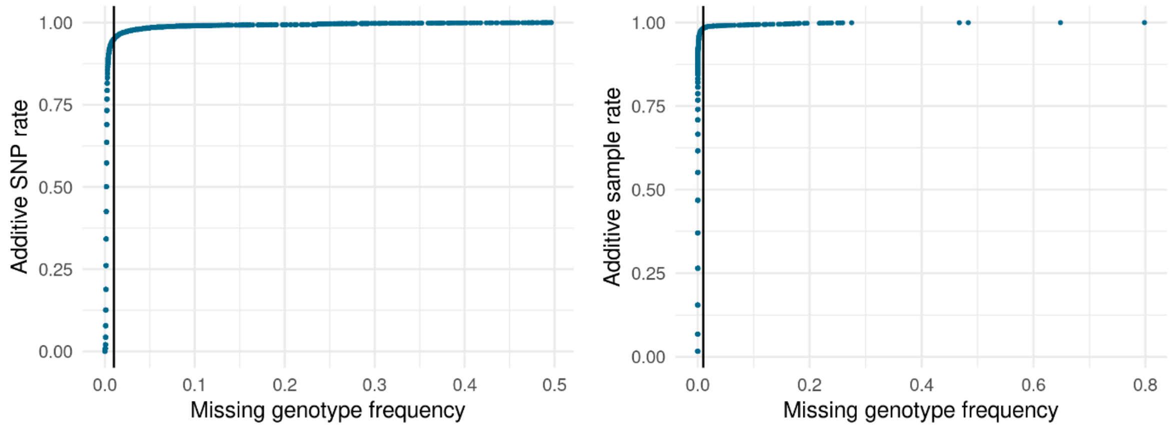


**Figure S3:** Additive SNP and sample call-rate cumulative distributions. Left: SNP call-rate; right: sample call-rate. Black line: chosen threshold=0.01.

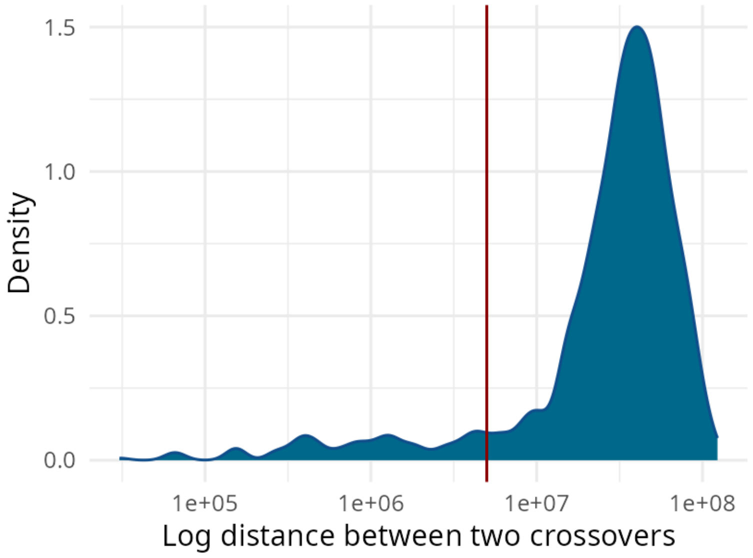


**Figure S4:** Distribution of the distance between two crossovers per chromosome per meiosis.

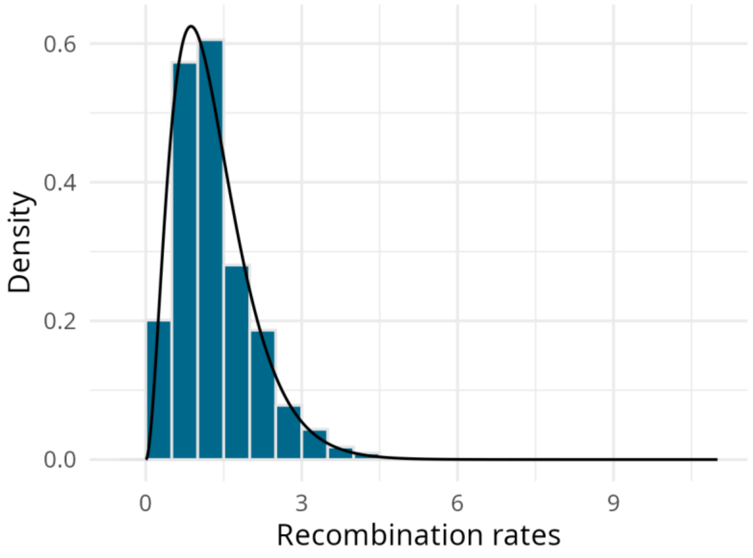


**Figure S5:** Density distribution of the Maximum Likelihood estimates of recombination rates on 1Mb intervals of the genome. Black: gamma law; extracted parameters α=3.08 & β=2.20.

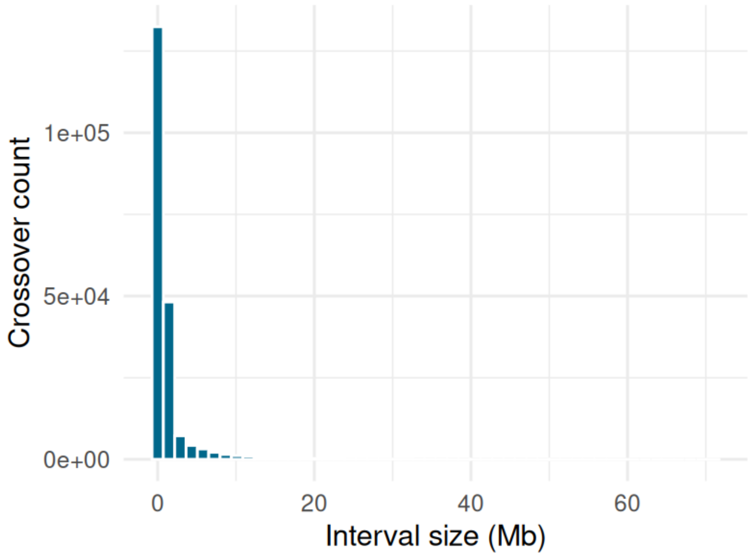


**Figure S6:** Distribution of crossover detection interval sizes.

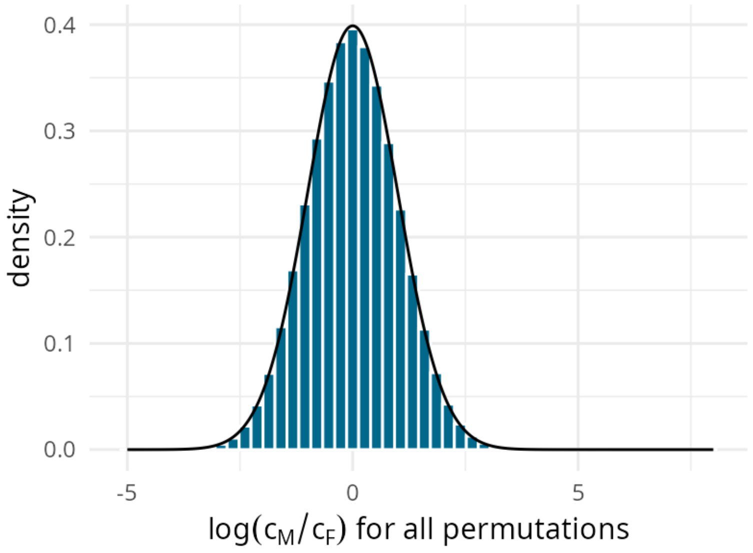


**Figure S7:** Distribution of all permuted recombination rate log ratio for all genomic intervals.

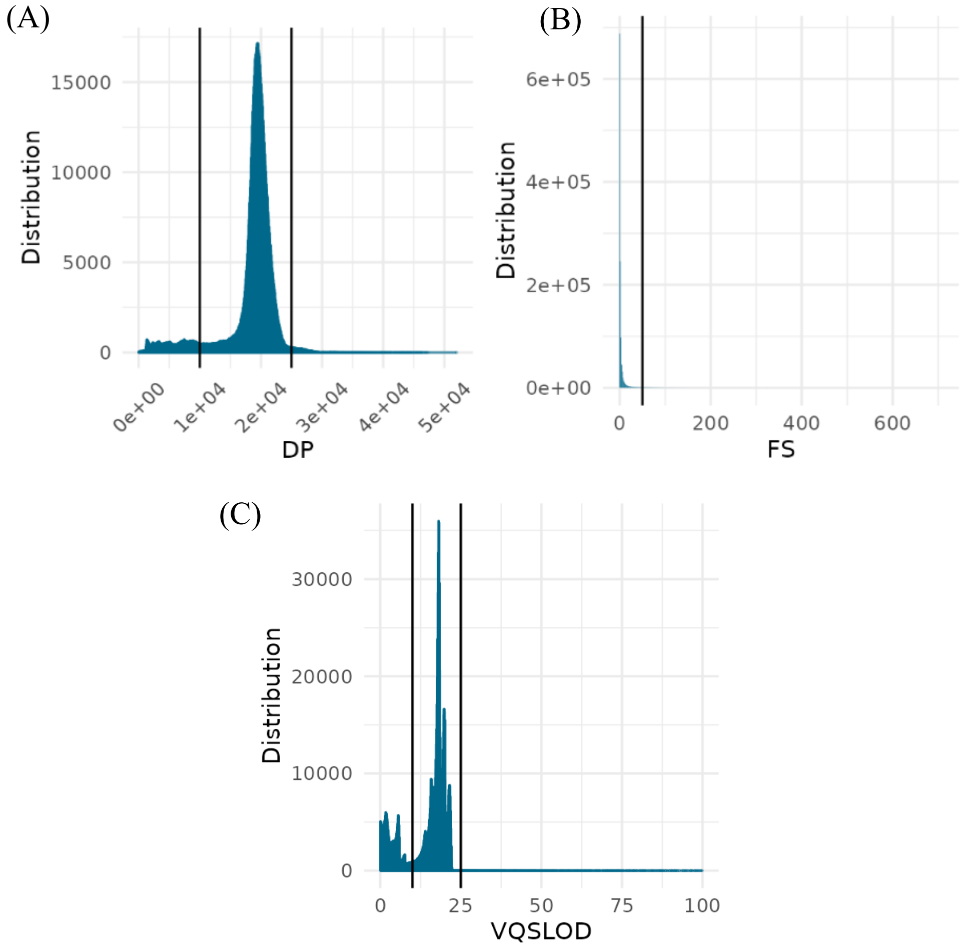

**Figure S8:** SNP distribution of sequencing and variant calling quality parameters. A: DP parameter; B: FS parameter; C: VQSLOD parameter.

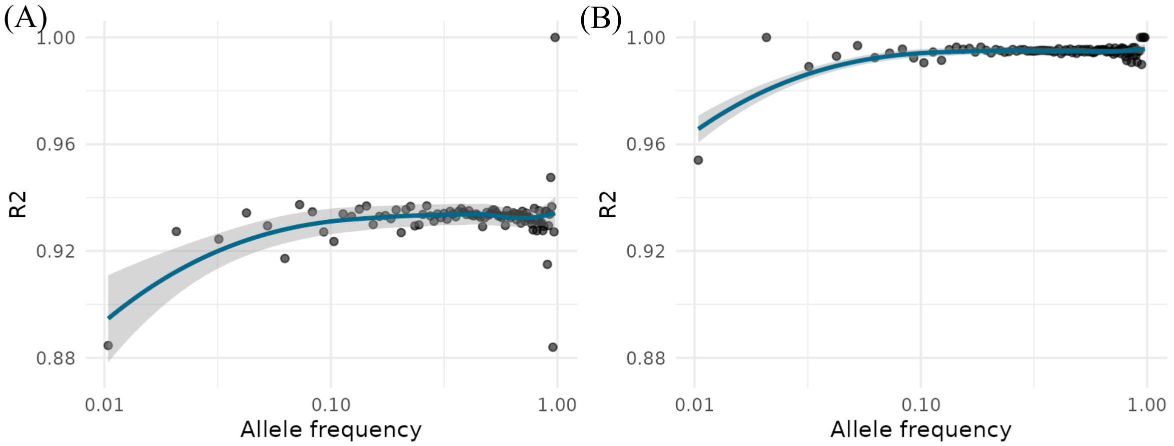


**Figure S9**: $R^2$ evolution according to allele frequency. (A) Before genotype imputation. Mean $R^2$=0.928. (B) After genotype imputation. Mean $R^2$=0.990.

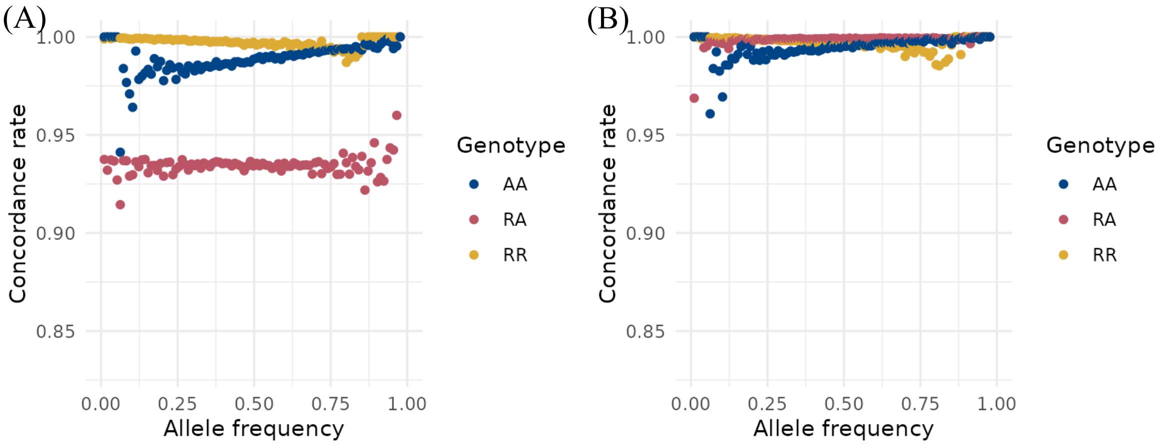


**Figure S10:** Genotype concordance according to allele frequency. (A) Before genotype imputation. Mean genotype concordance=0.974. (B) After genotype imputation. Mean genotype concordance=0.997.

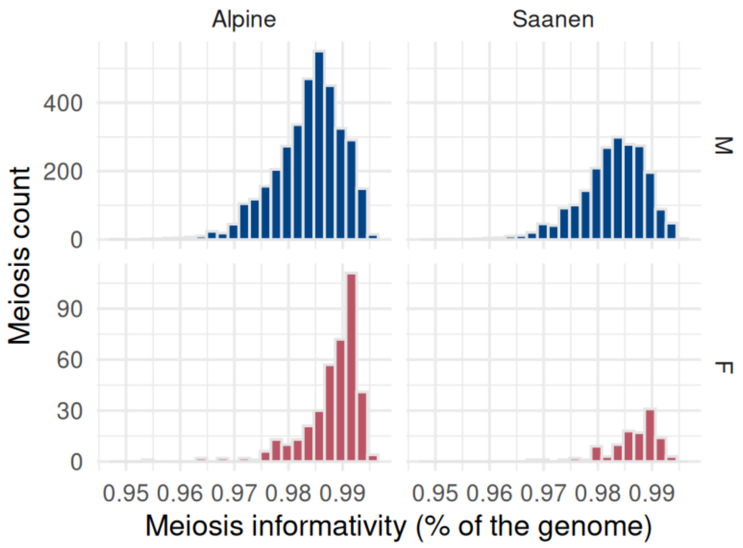


**Figure S11:** Distribution the proportion of the autosomal genome informative for crossovers in parents (M=males vs F=females).

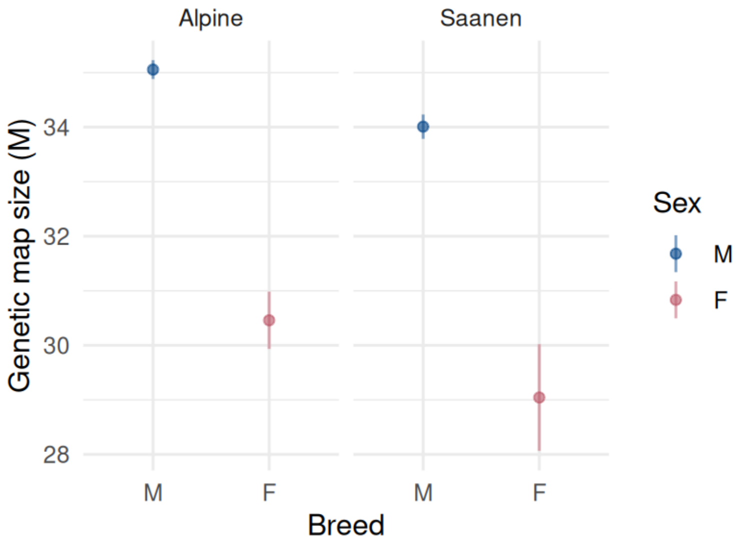


**Figure S12:** Comparison of recombination map size estimates between populations. Confidence interval = $\mu \pm 2 \times standard\ error$.

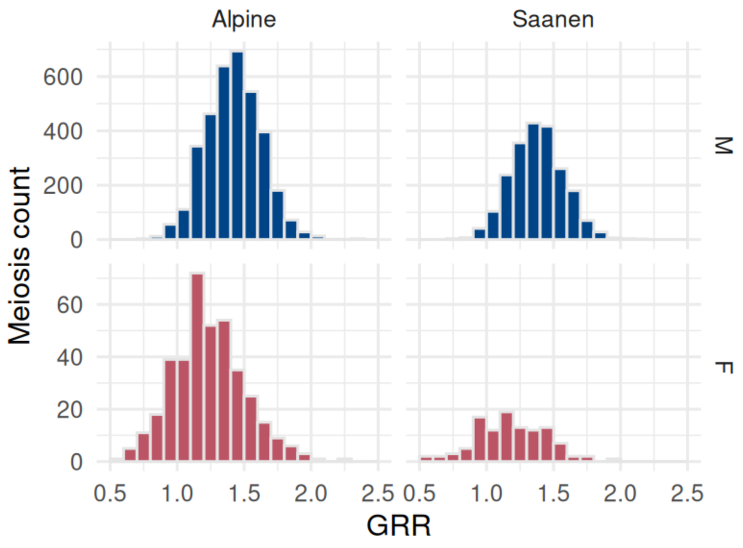


**Figure S13:** Distribution of genome-wide recombination rates (GRR) in meioses.

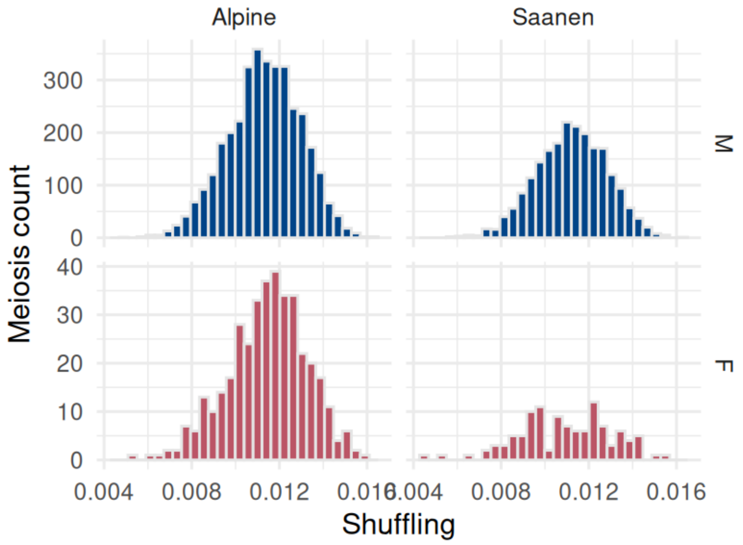


**Figure S14:** Distribution of intra-chromosomic shuffling in meioses.

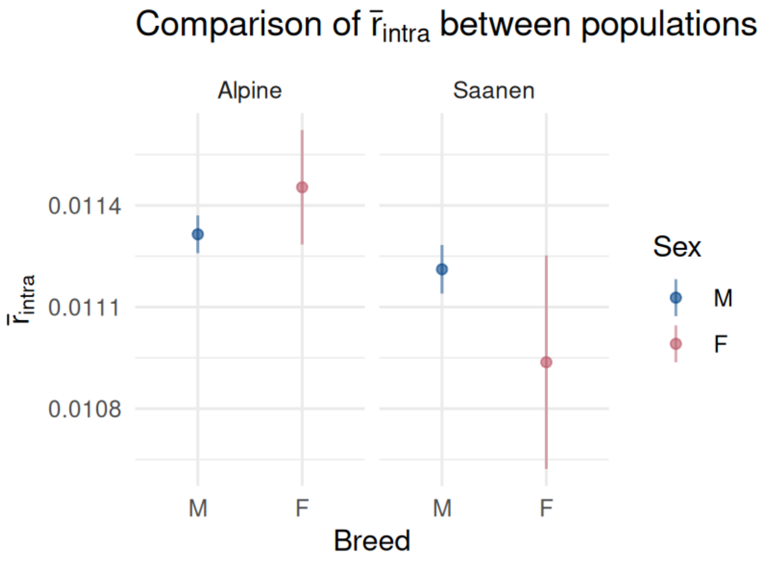


**Figure S15**: Comparison of intra-chromosomic shuffling estimates between populations. Confidence interval = $\mu \pm 2 \times standard\ error$.

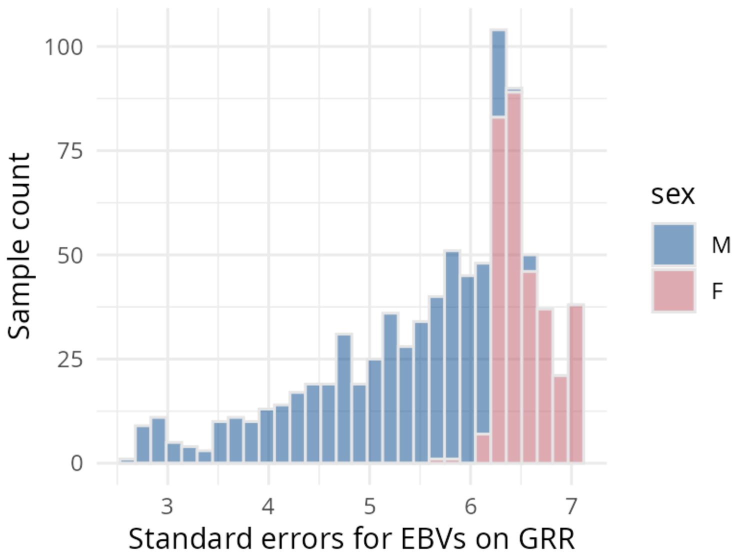


**Figure S16:** Distribution of standard errors for EBVs on GRR.

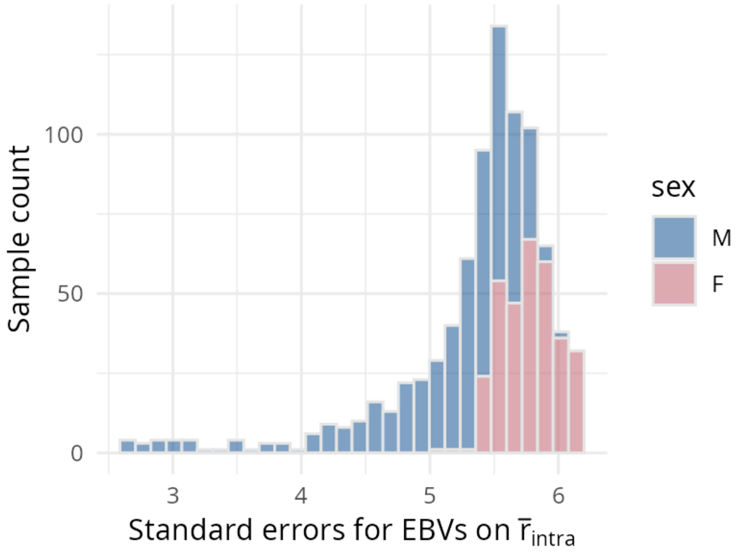


**Figure S17:** Distribution of standard errors for EBVs on $\bar{r}_{intra}$
