## Additional File 5 for "Genome landscape and genetic architecture of recombination in domestic goats (*Capra Hircus*)"

### Chromosome-specific recombination maps

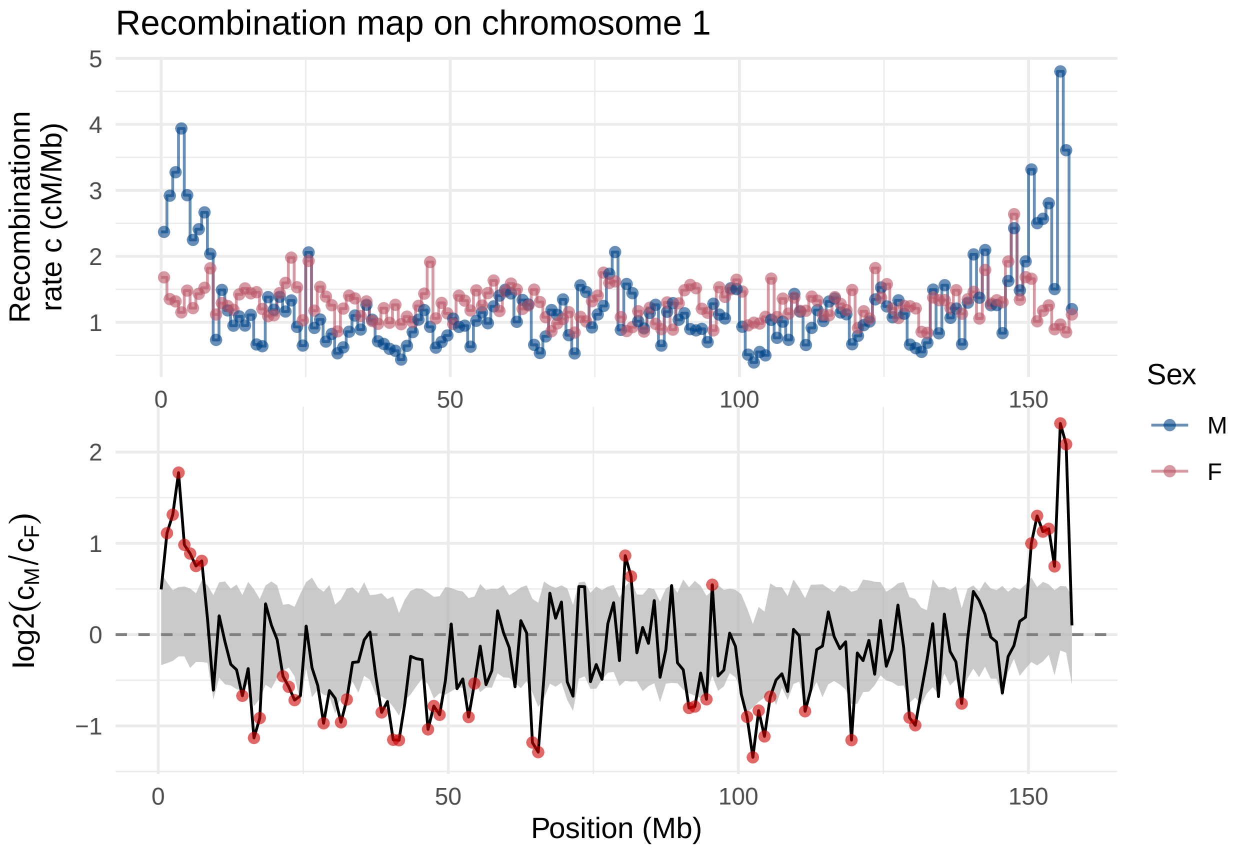

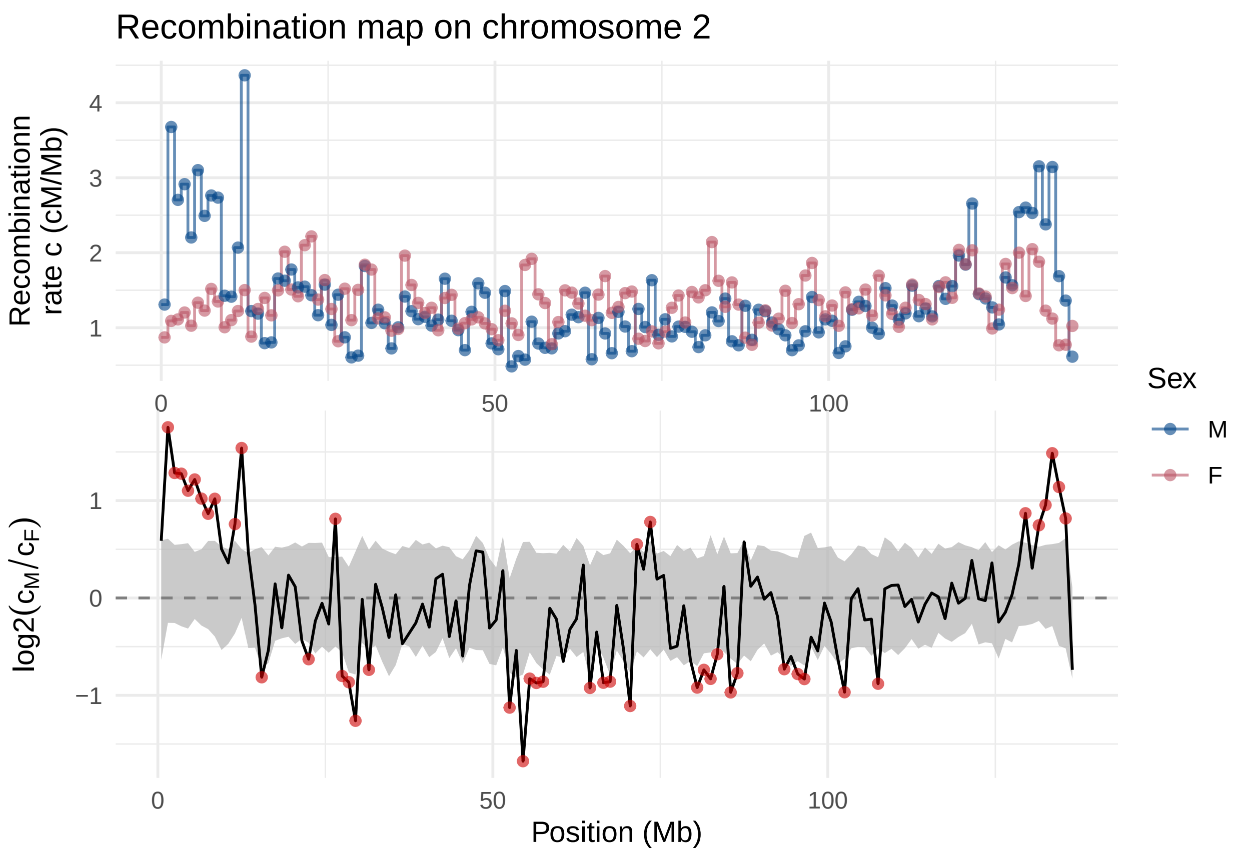

Recombination map on chromosome 2
Recombinationn
rate c (cM/Mb)
4
3
2
1
0
50
100
Sex
M
F
log2($c_M/c_F$)
1
0
−1
0
50
100
Position (Mb)

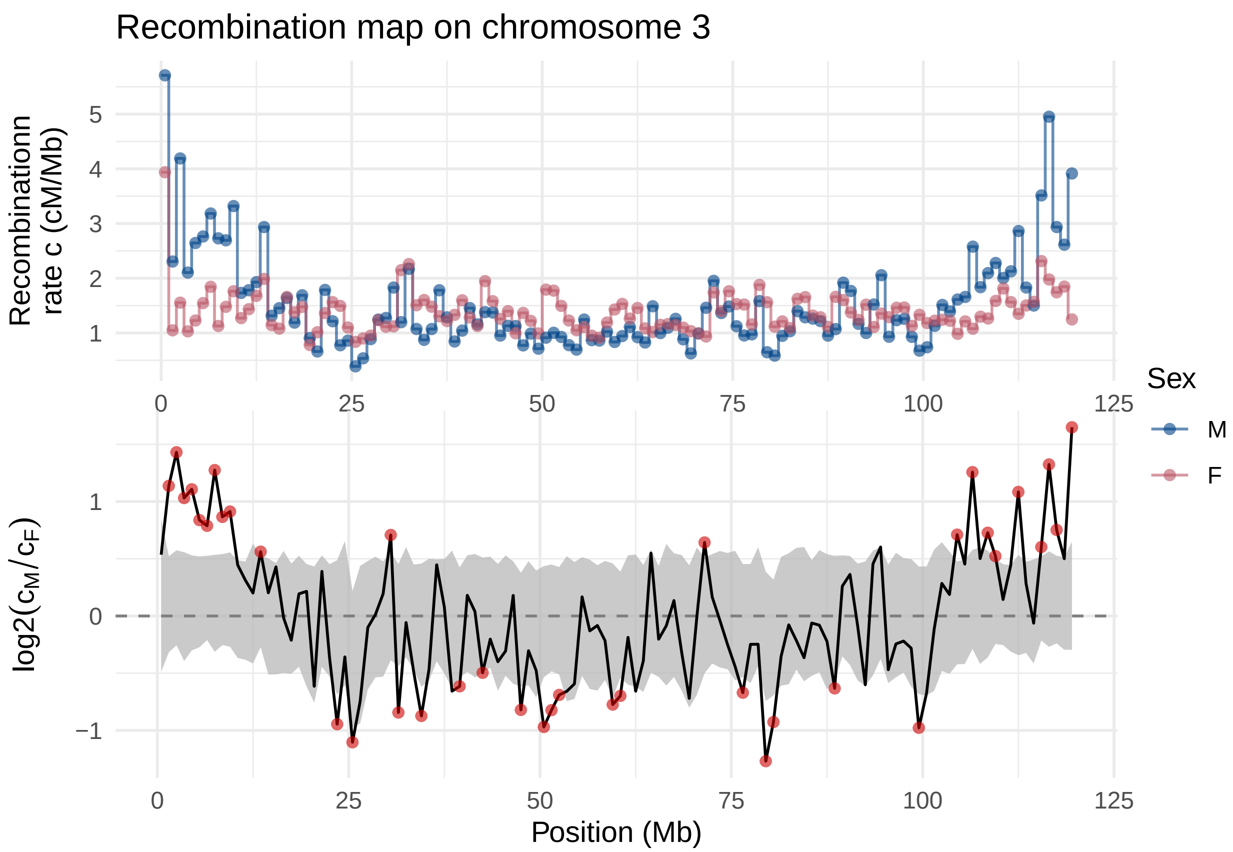

Recombination map on chromosome 3
Recombinationn rate c (cM/Mb)
5
4
3
2
1
0
25
50
75
100
125
Sex
M
F
log2(c_M/c_F)
1
0
−1
0
25
50
75
100
125
Position (Mb)

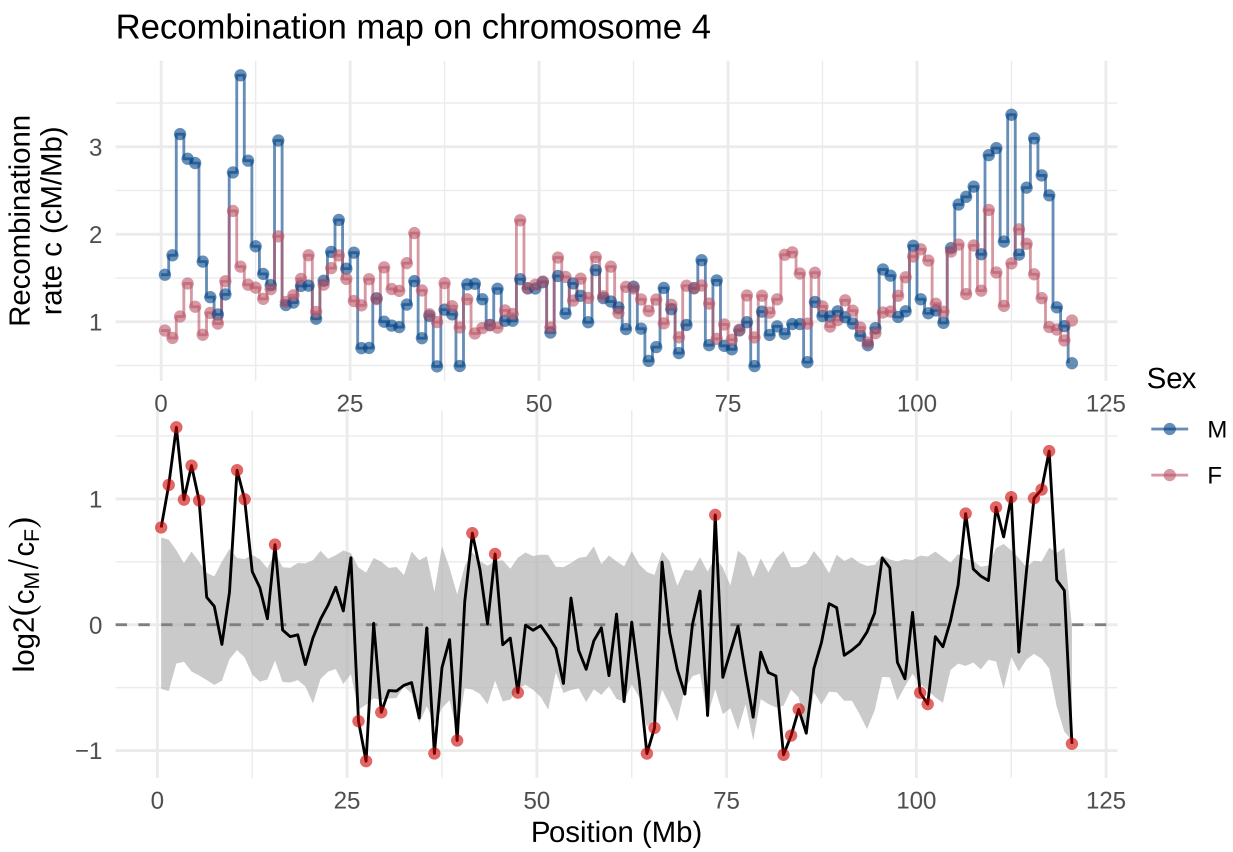

Recombination map on chromosome 4
Recombinationn rate c (cM/Mb)
3
2
1
0
25
50
75
100
125
Sex
M
F
log2(c_M/c_F)
1
0
−1
Position (Mb)

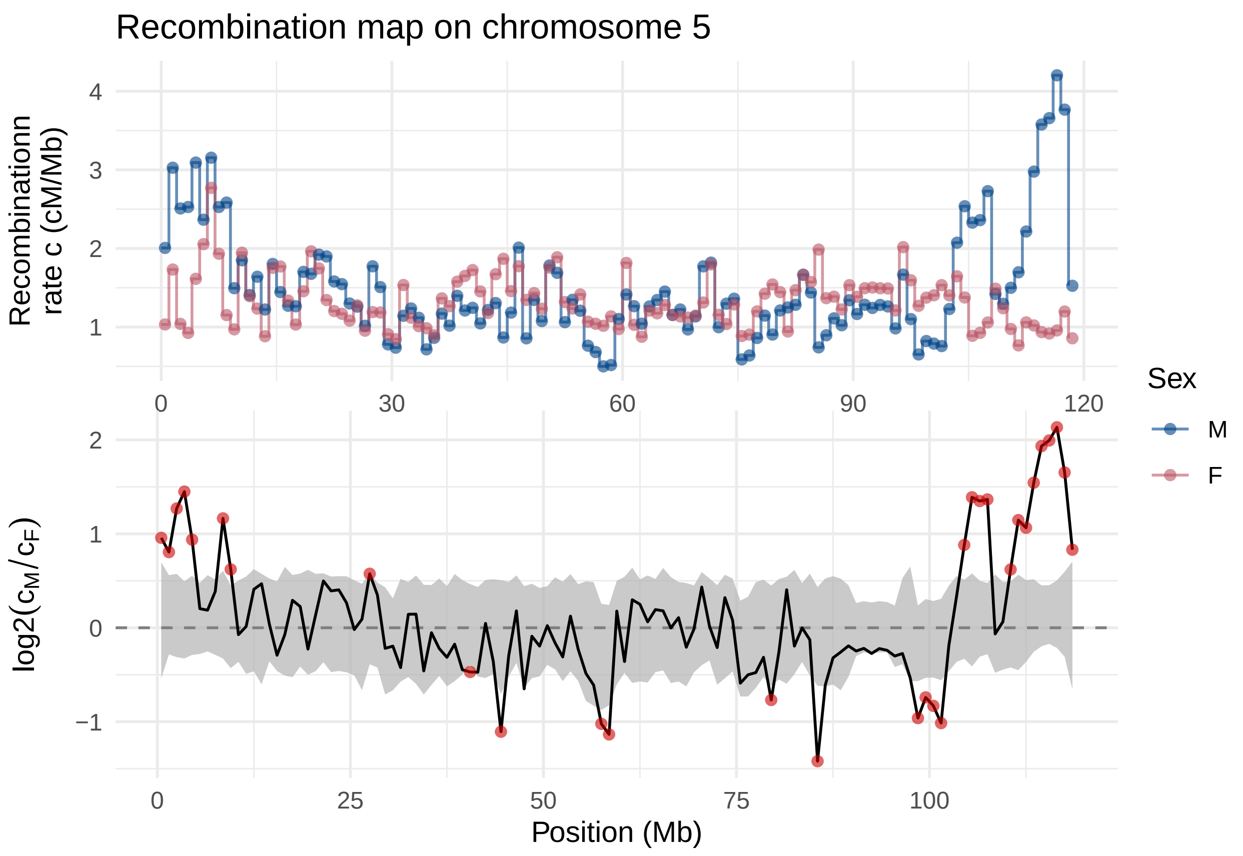
Recombination map on chromosome 5
Recombinationn rate c (cM/Mb)
4
3
2
1
0
30
60
90
120
Sex
M
F
log2(cM/cF)
2
1
0
−1
0
25
50
75
100
Position (Mb)

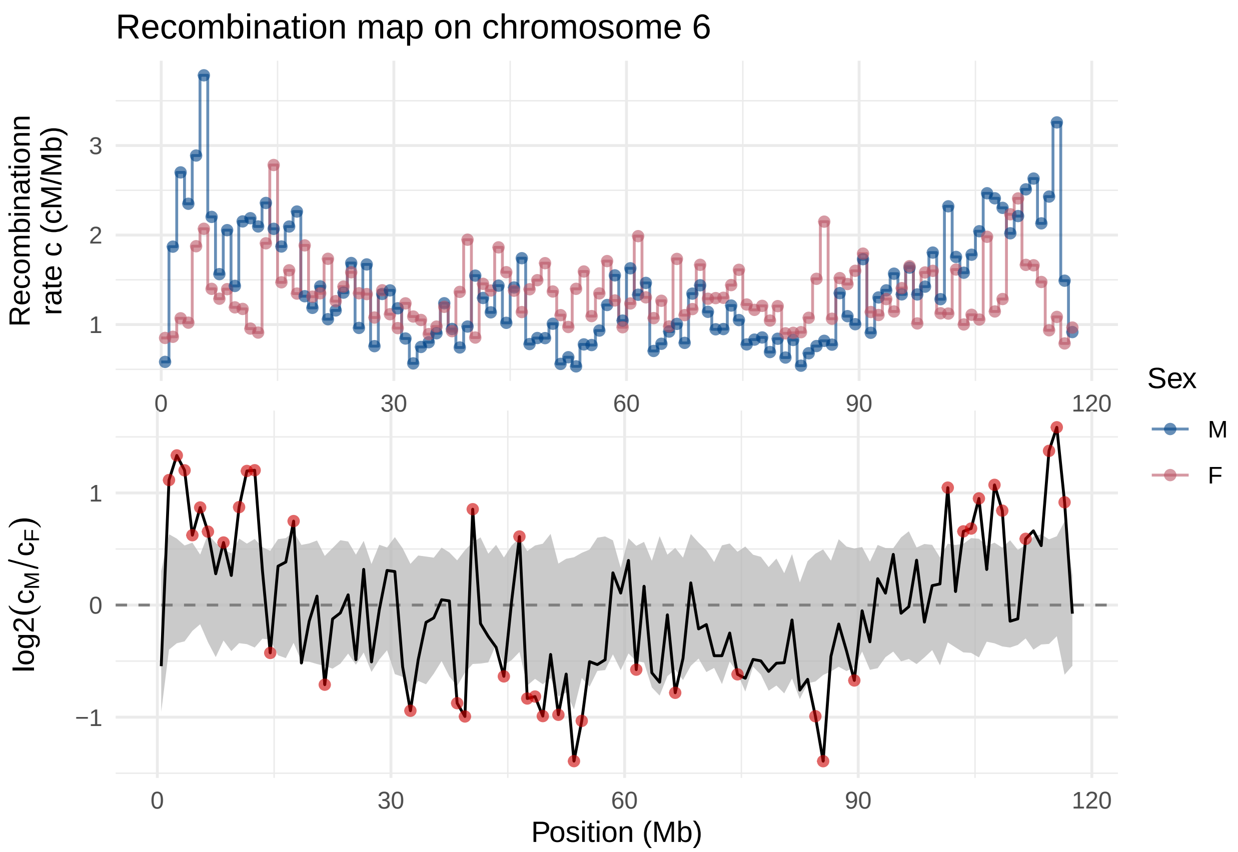

Recombination map on chromosome 6
Recombinationn rate c (cM/Mb)
3
2
1
0
30
60
90
120
Sex
M
F
log2(c_M/c_F)
1
0
−1
0
30
60
90
120
Position (Mb)

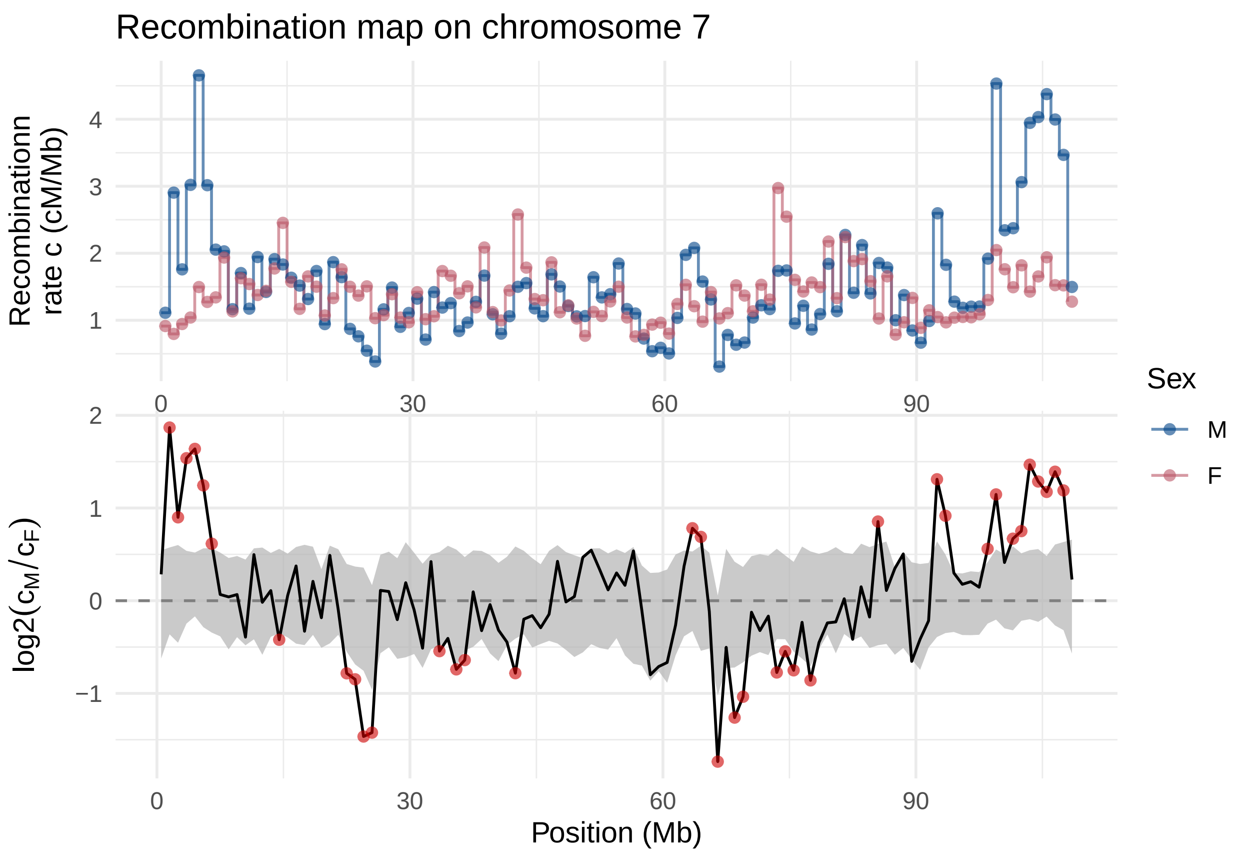

Recombination map on chromosome 7
Recombinationn
rate c (cM/Mb)
log2(c_M/c_F)
Position (Mb)
Sex
M
F

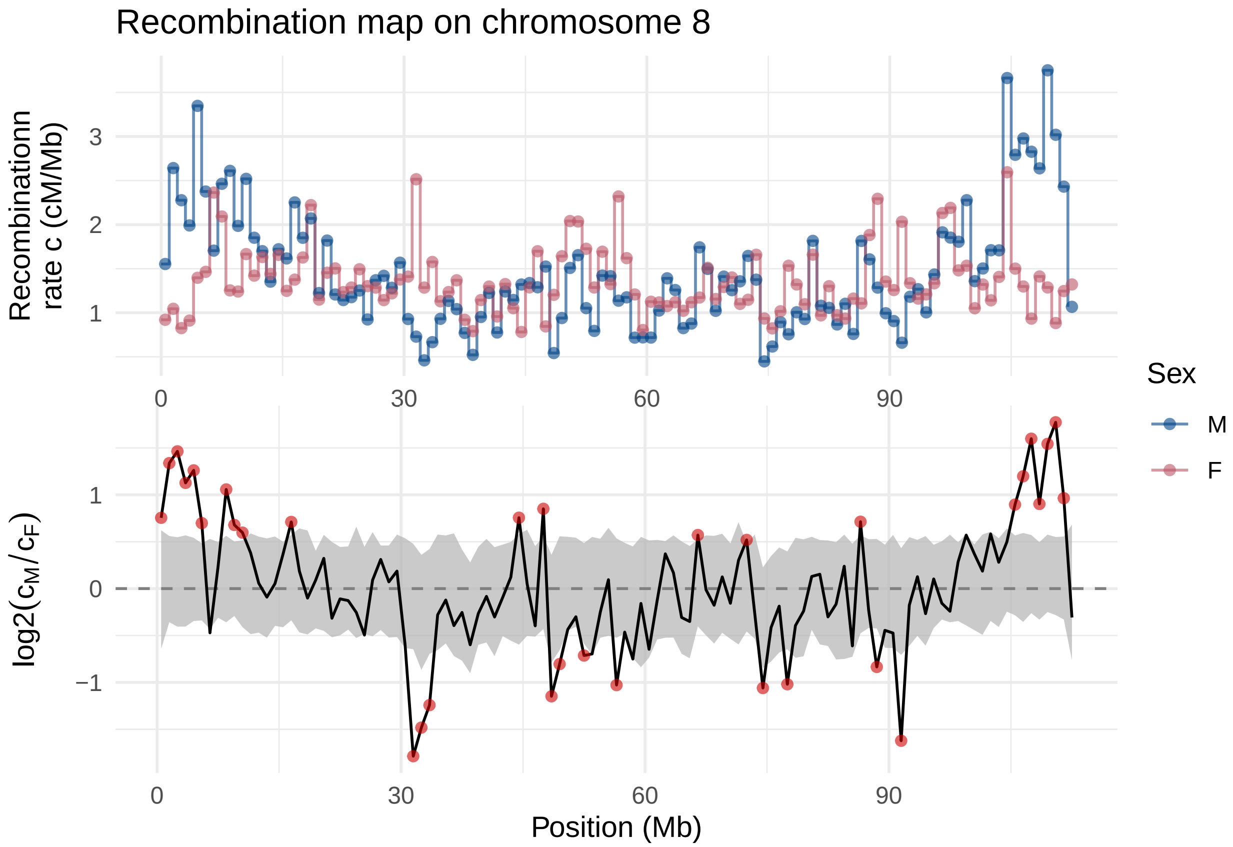

Recombination map on chromosome 8
Recombinationn rate c (cM/Mb)
3
2
1
0
30
60
90
Sex
M
F
log2(c_M/c_F)
1
0
−1
0
30
60
90
Position (Mb)

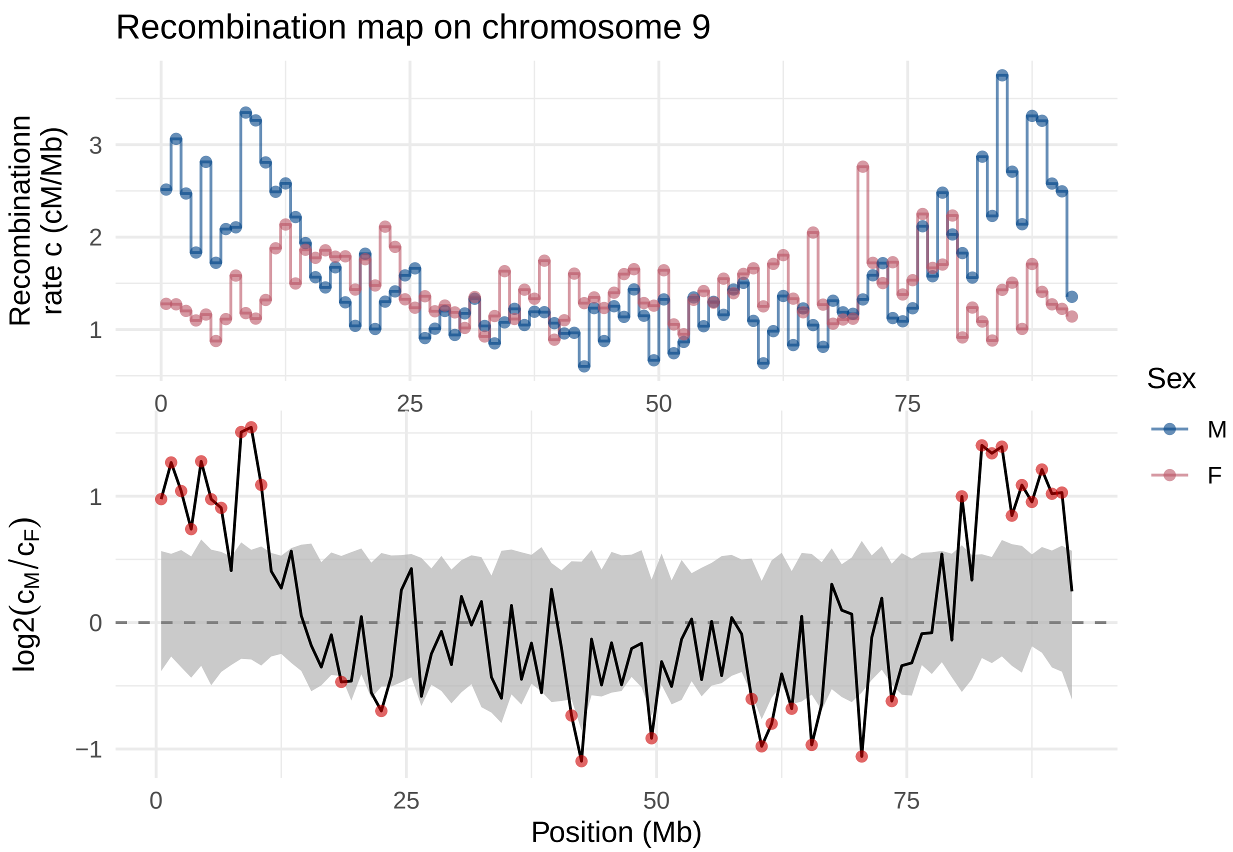
Recombination map on chromosome 9
Recombinationn
rate c (cM/Mb)
3
2
1
0
25
50
75
Sex
M
F
log2(c_M/c_F)
1
0
−1
0
25
50
75
Position (Mb)

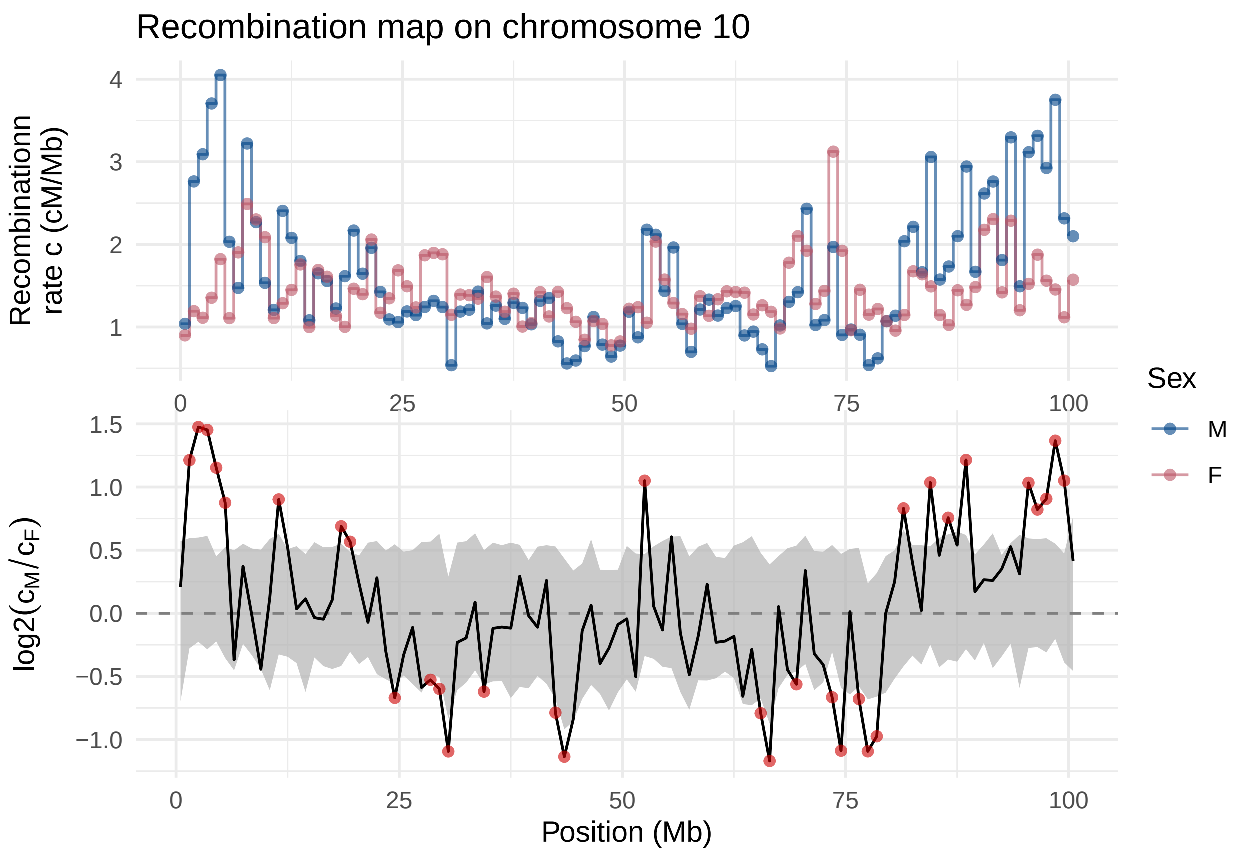

Recombination map on chromosome 10
Recombinationn rate c (cM/Mb)
log2(cM/cF)
Position (Mb)
Sex
M
F

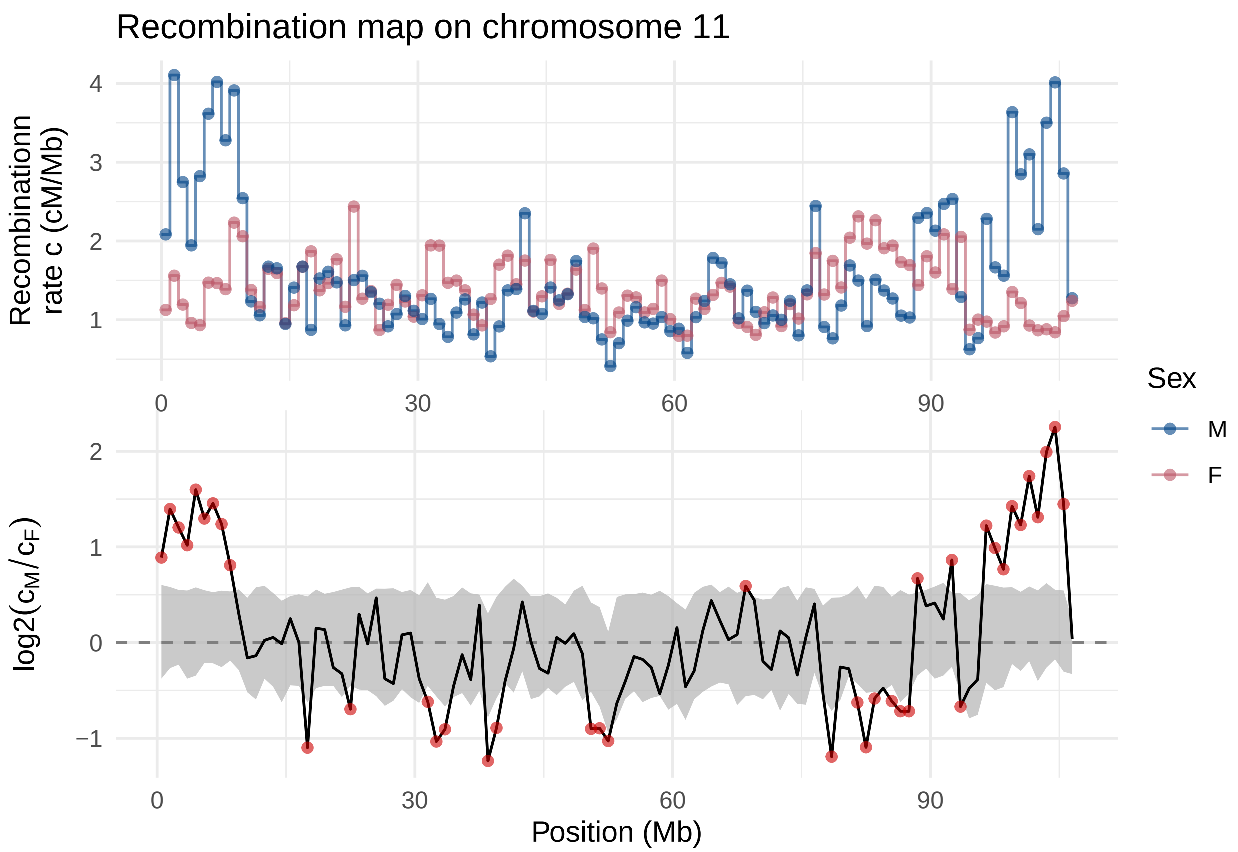
Recombination map on chromosome 11
Recombinationn rate c (cM/Mb)
4
3
2
1
0
30
60
90
Sex
M
F
log2(cM/cF)
2
1
0
−1
0
30
60
90
Position (Mb)

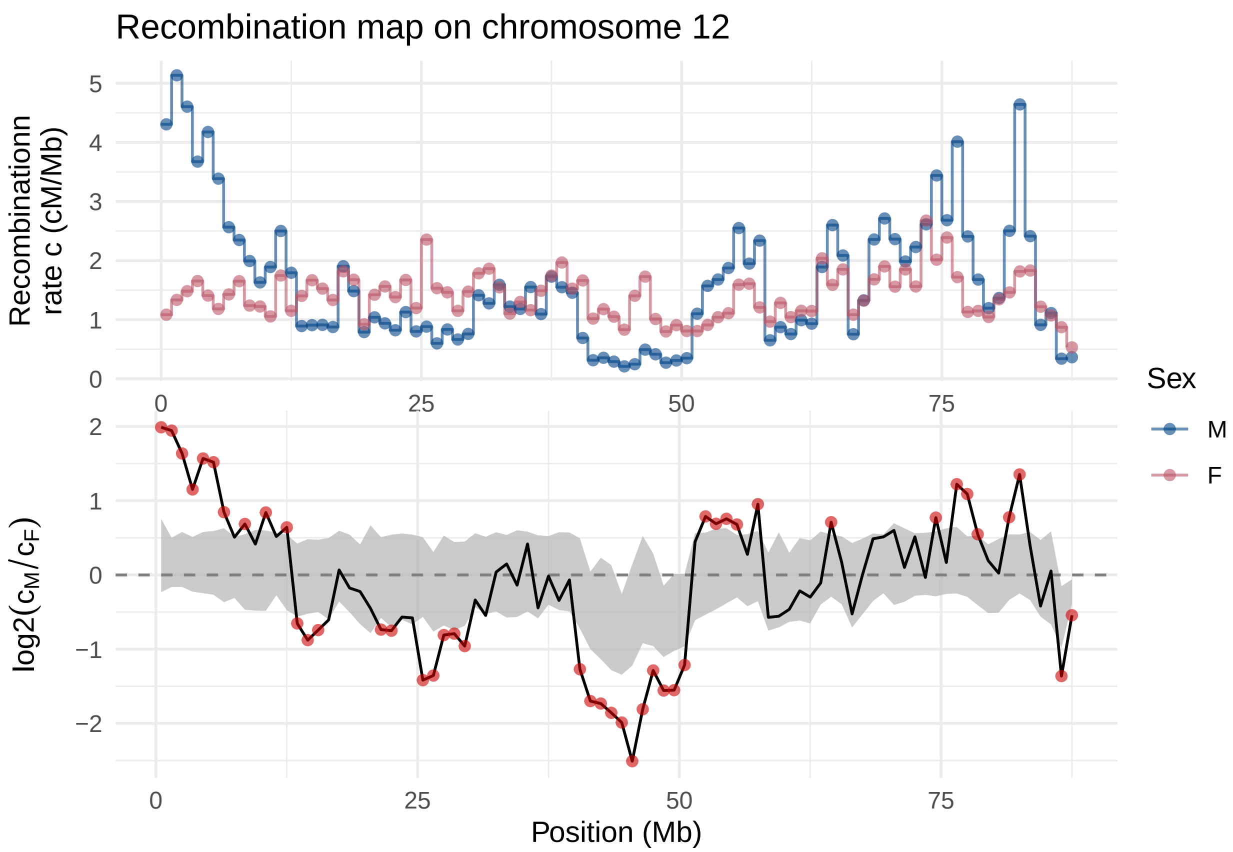

Recombination map on chromosome 12
Recombinationn rate c (cM/Mb)
5
4
3
2
1
0
0
25
50
75
Sex
M
F
$\log2(c_M/c_F)$
2
1
0
−1
−2
0
25
50
75
Position (Mb)

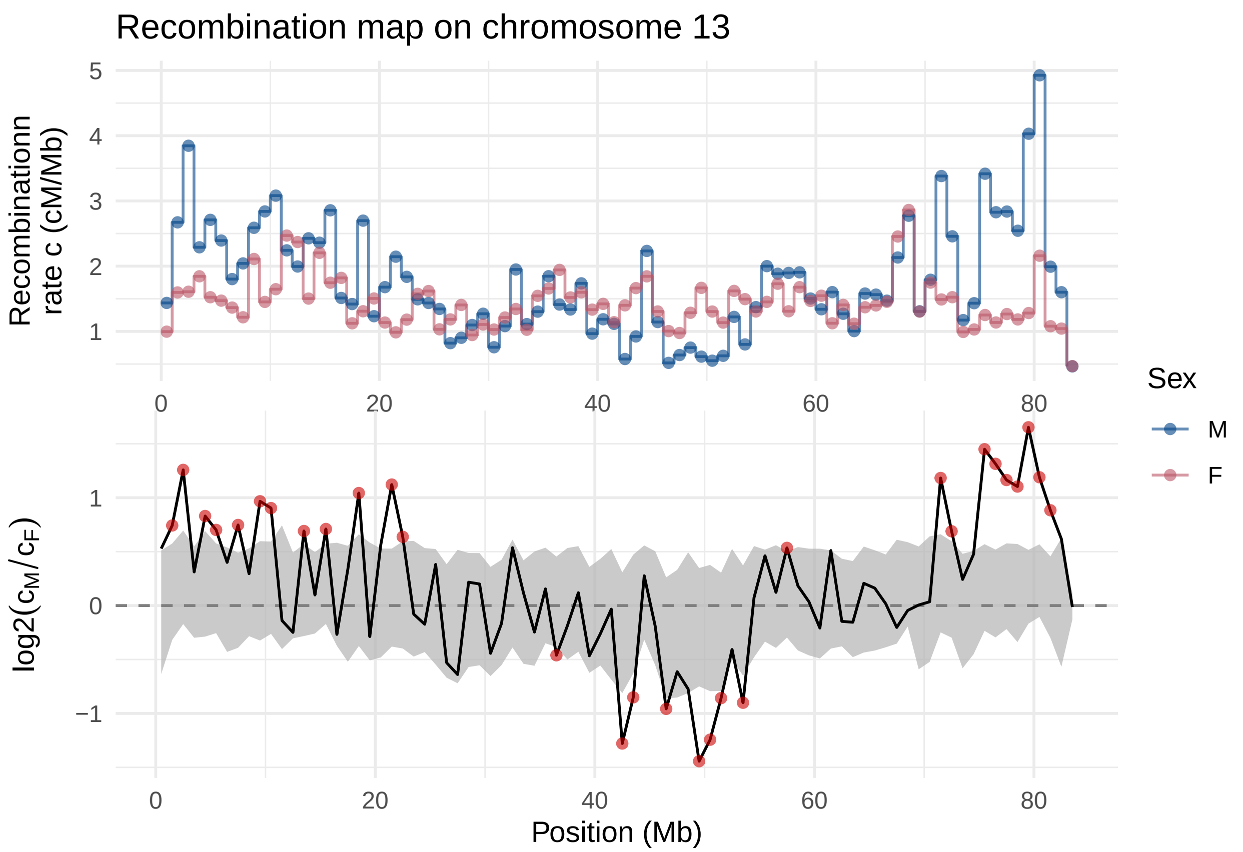
Recombination map on chromosome 13
Recombinationn
rate c (cM/Mb)
log2(c_M/c_F)
Position (Mb)
Sex
M
F

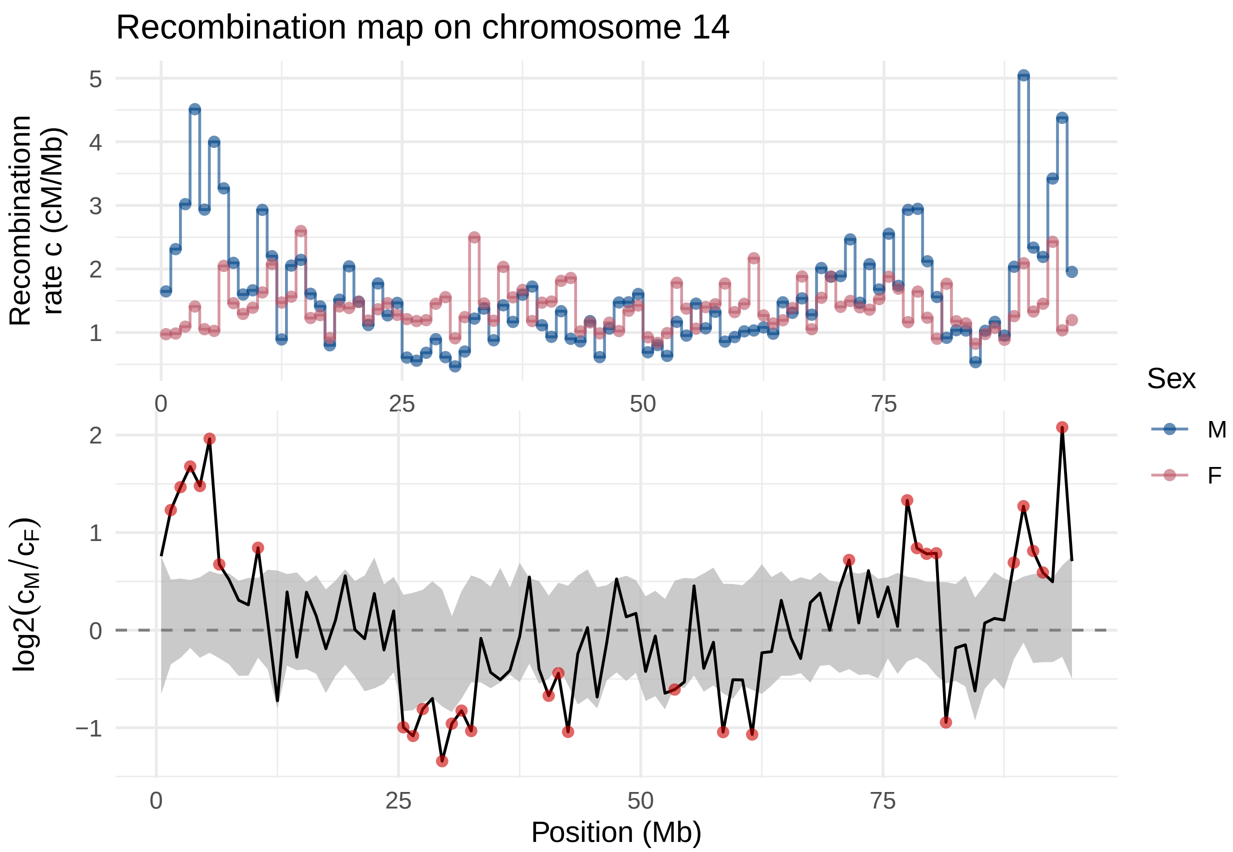

Recombination map on chromosome 14
Recombinationn rate c (cM/Mb)
5
4
3
2
1
0
25
50
75
Sex
M
F
log2(cM/cF)
2
1
0
−1
Position (Mb)

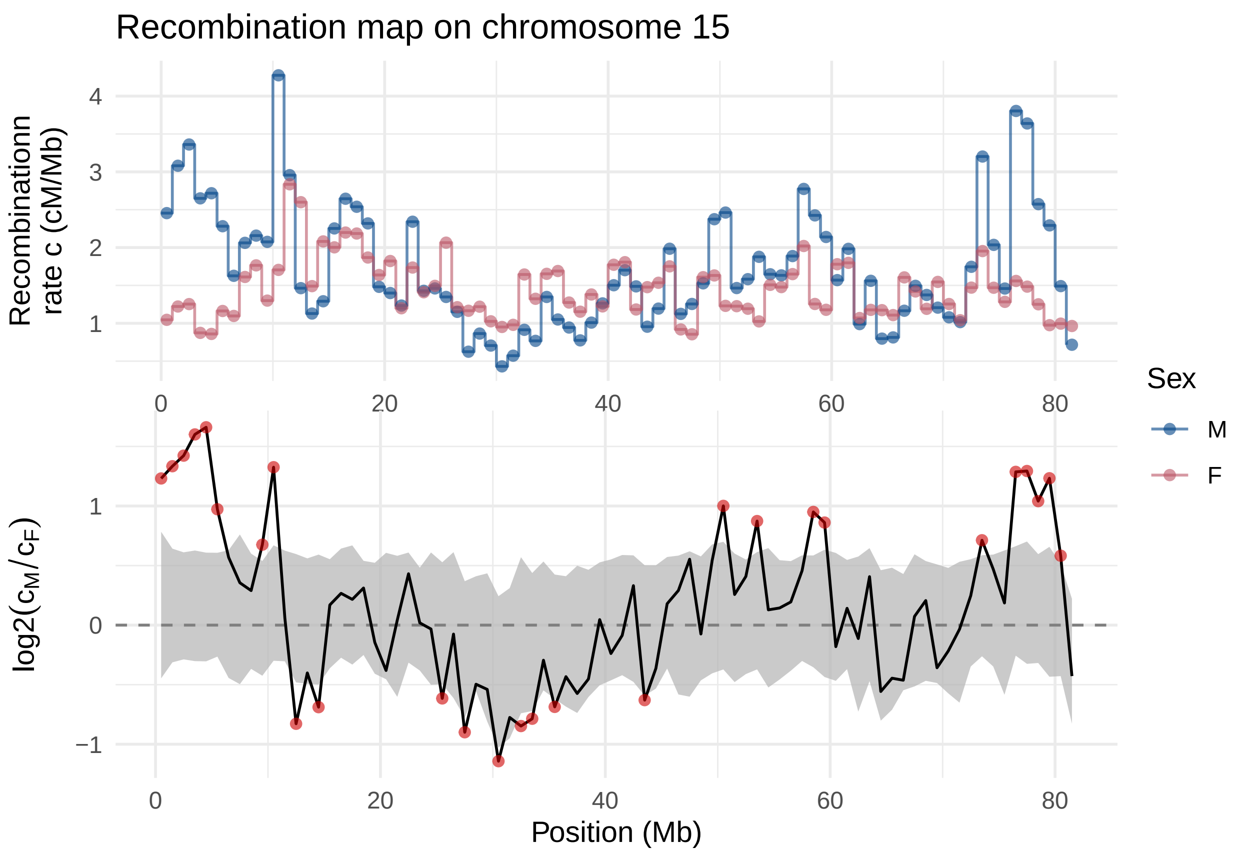

Recombination map on chromosome 15
Recombinationn rate c (cM/Mb)
4
3
2
1
0
20
40
60
80
log2(c_M/c_F)
1
0
−1
0
20
40
60
80
Position (Mb)
Sex
M
F

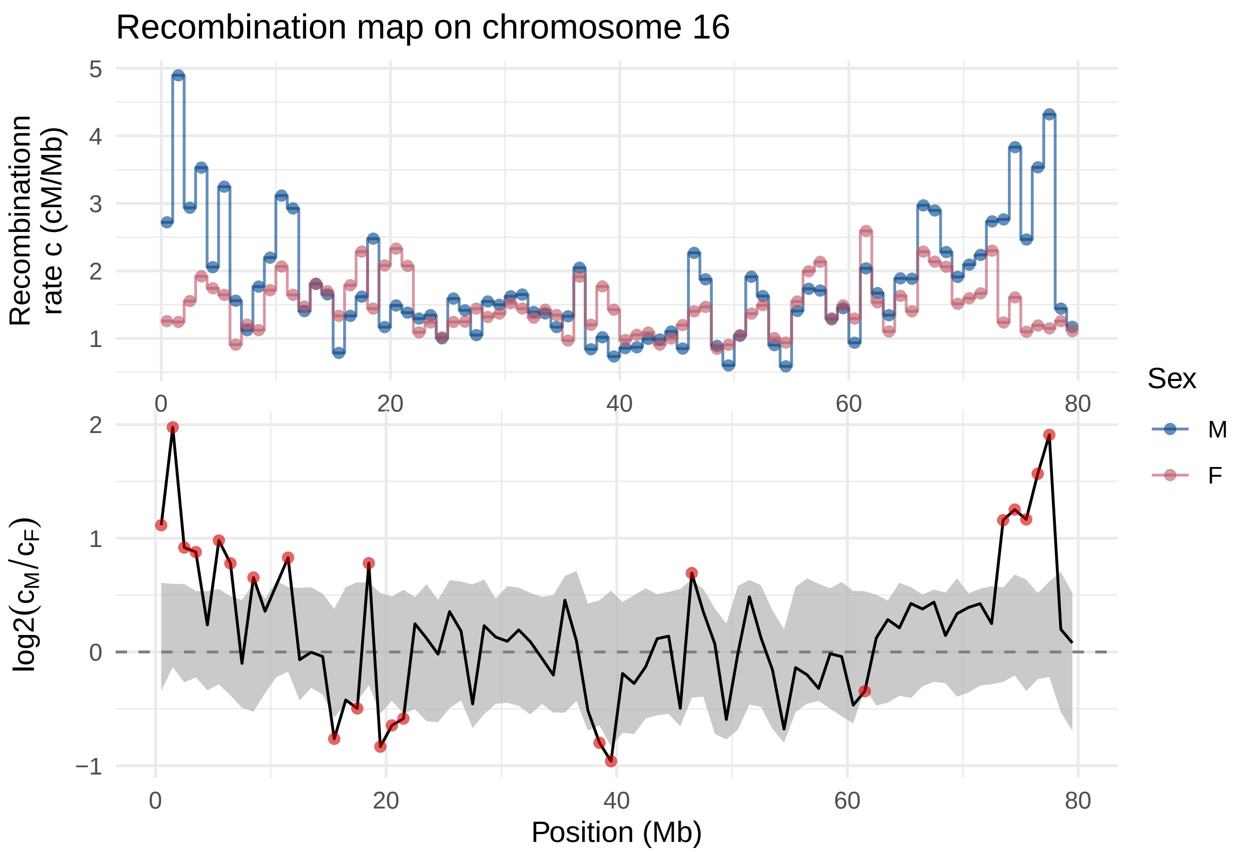
Recombination map on chromosome 16
Recombinationn rate c (cM/Mb)
log2(cM/cF)
Position (Mb)
Sex
M
F
0
20
40
60
80
1
2
3
4
5
−1

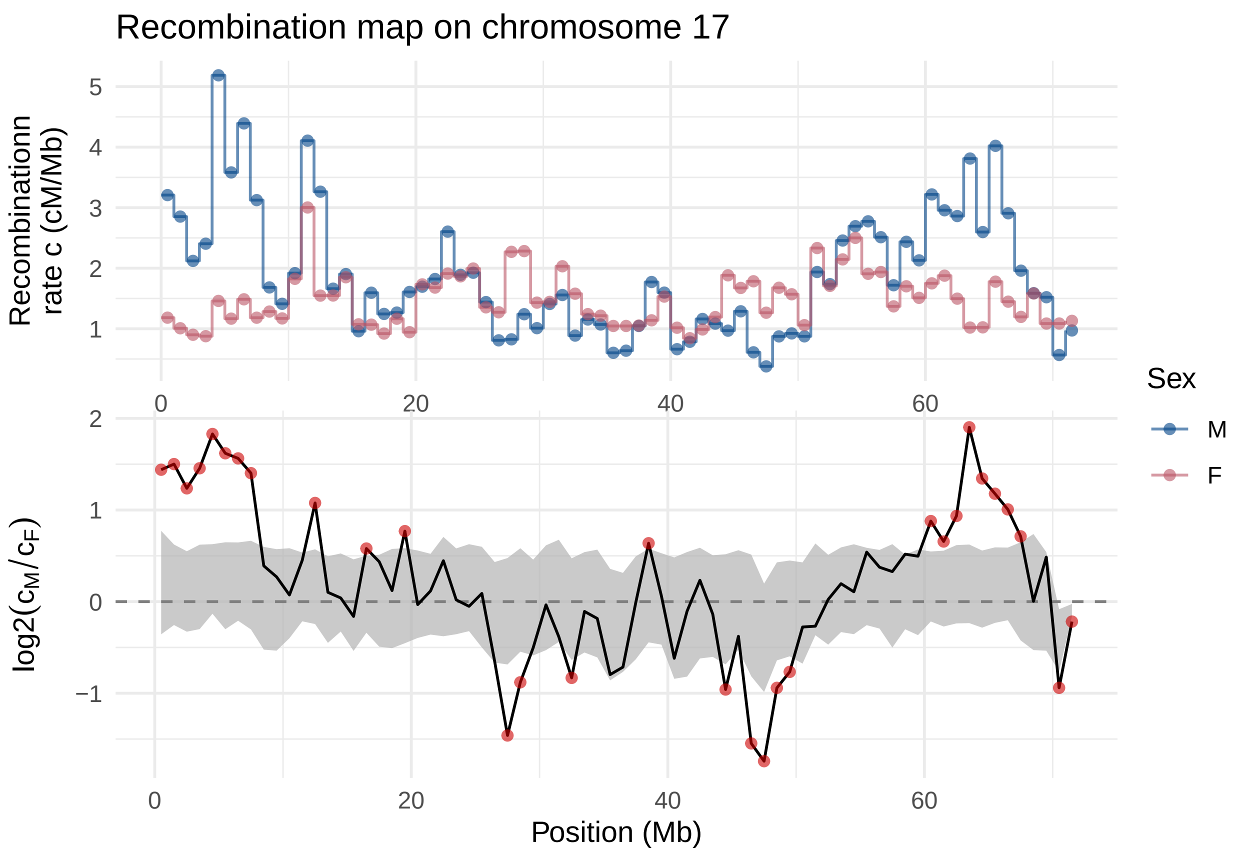

Recombination map on chromosome 17
Recombinationn rate c (cM/Mb)
5
4
3
2
1
0
20
40
60
Sex
M
F
log2(cM/cF)
2
1
0
−1
0
20
40
60
Position (Mb)

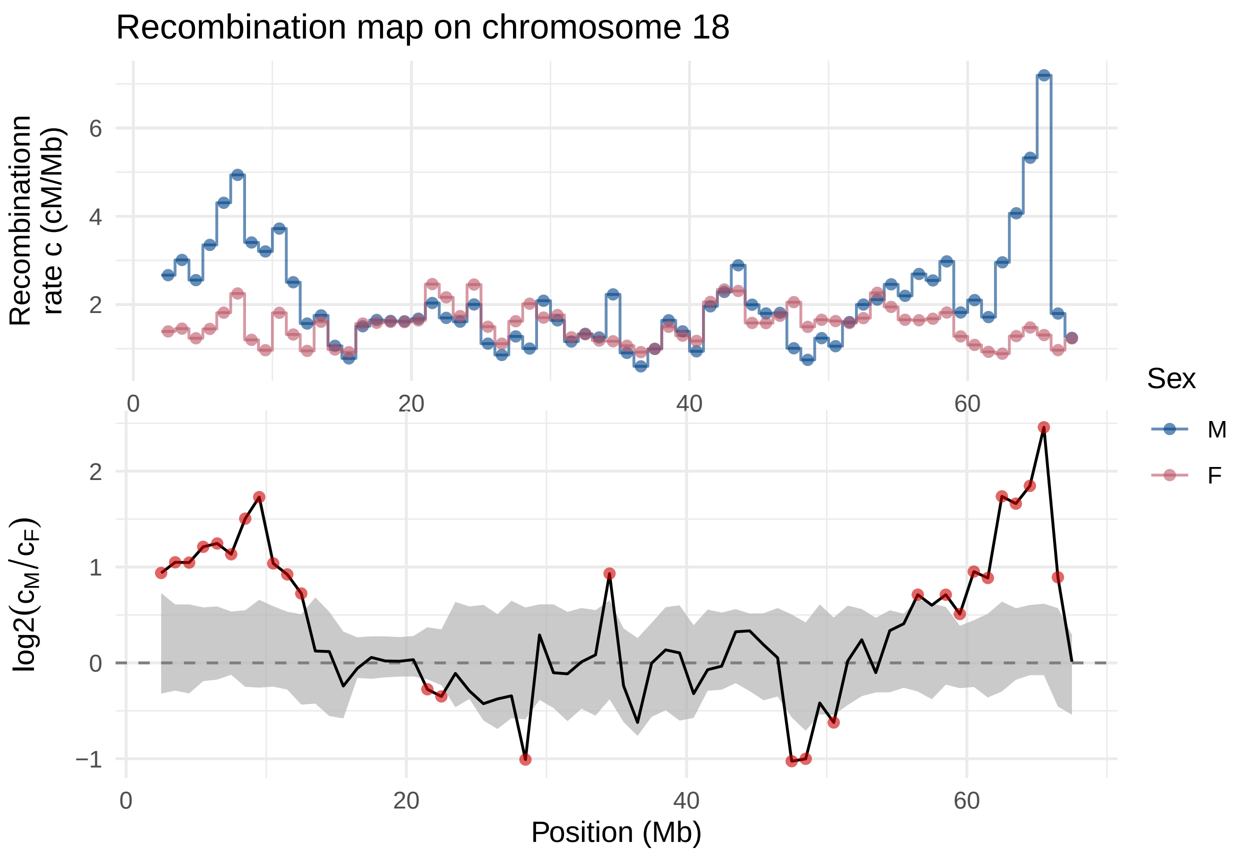

Recombination map on chromosome 18
Recombinationn rate c (cM/Mb)
6
4
2
0
20
40
60
Sex
M
F
log2(cM/cF)
2
1
0
−1
0
20
40
60
Position (Mb)

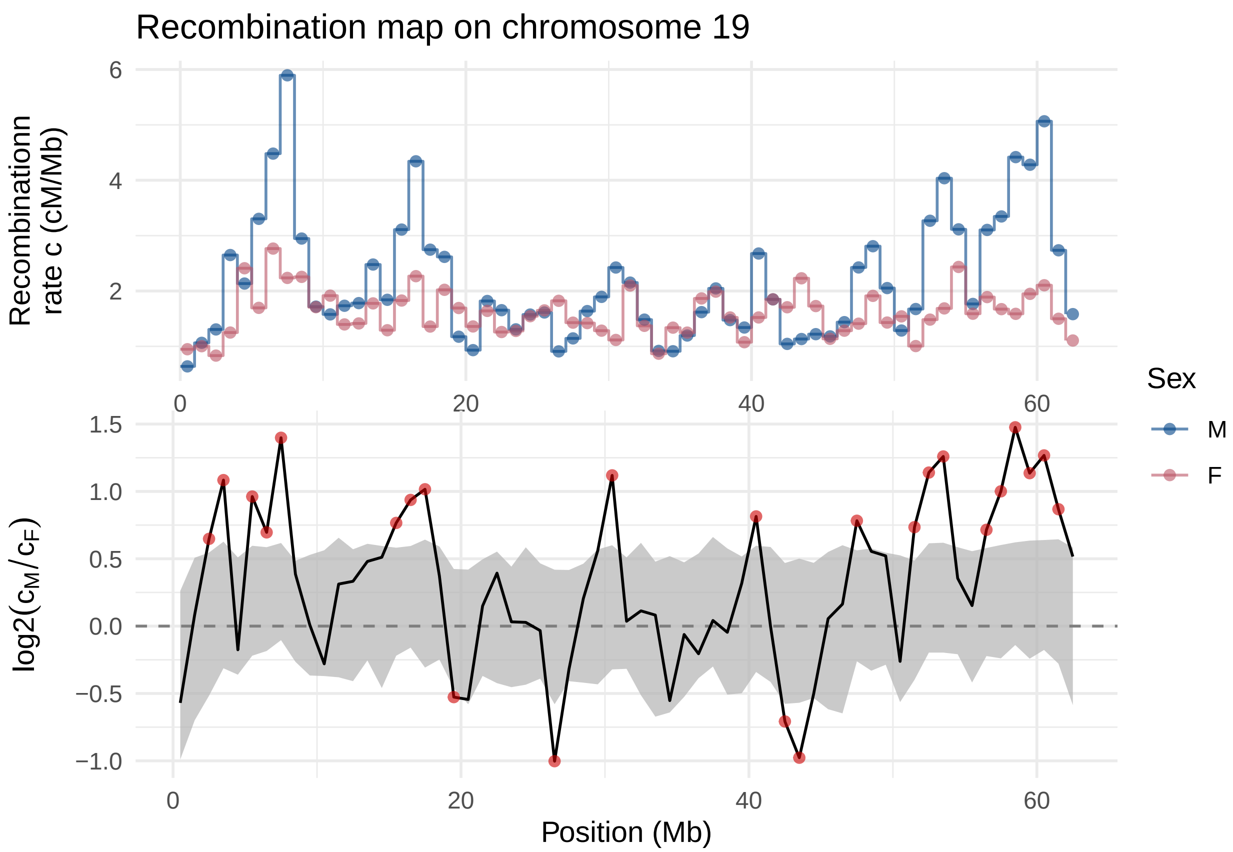
Recombination map on chromosome 19
Recombinationn rate c (cM/Mb)
6
4
2
0
20
40
60
Sex
M
F
log2(cM/cF)
1.5
1.0
0.5
0.0
−0.5
−1.0
0
20
40
60
Position (Mb)

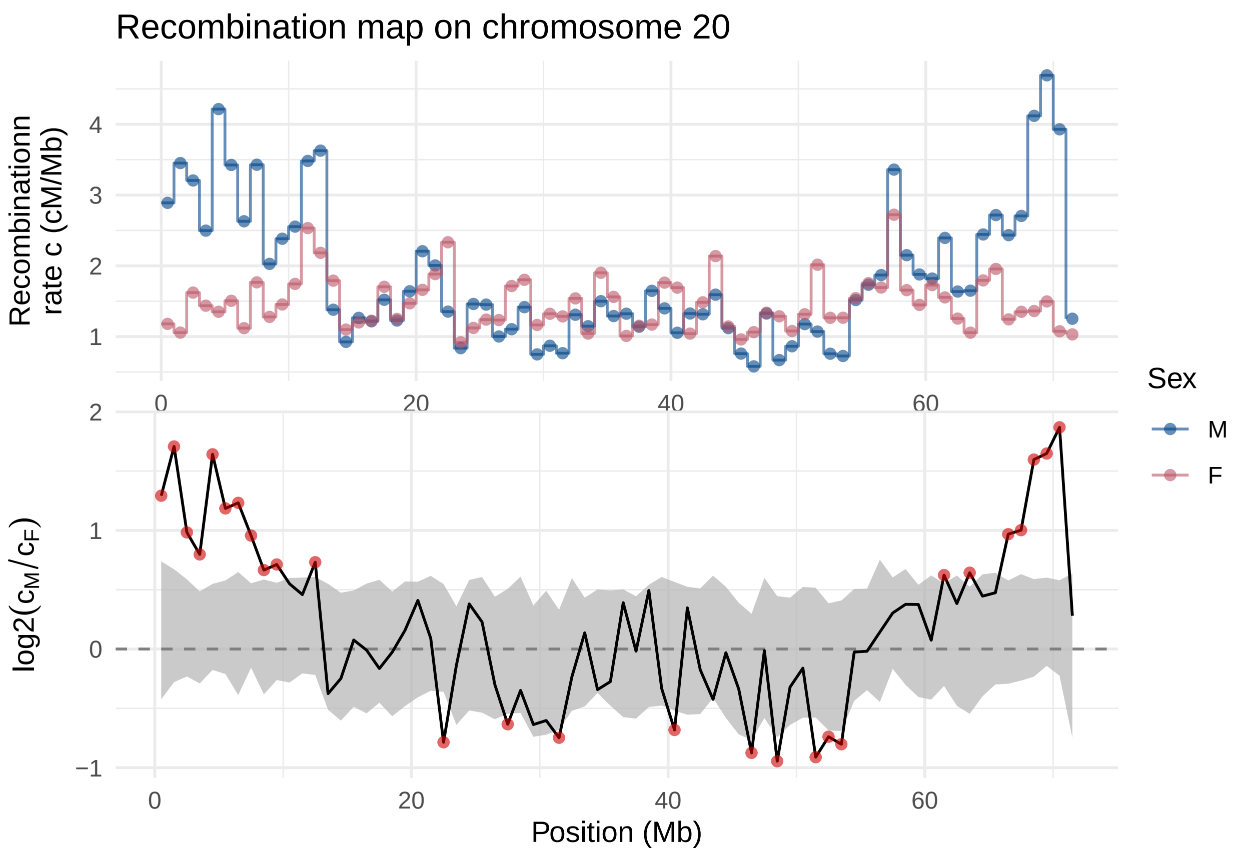

Recombination map on chromosome 20
Recombinationn rate c (cM/Mb)
4
3
2
1
0
20
40
60
Sex
M
F
log2(c_M/c_F)
2
1
0
−1
0
20
40
60
Position (Mb)

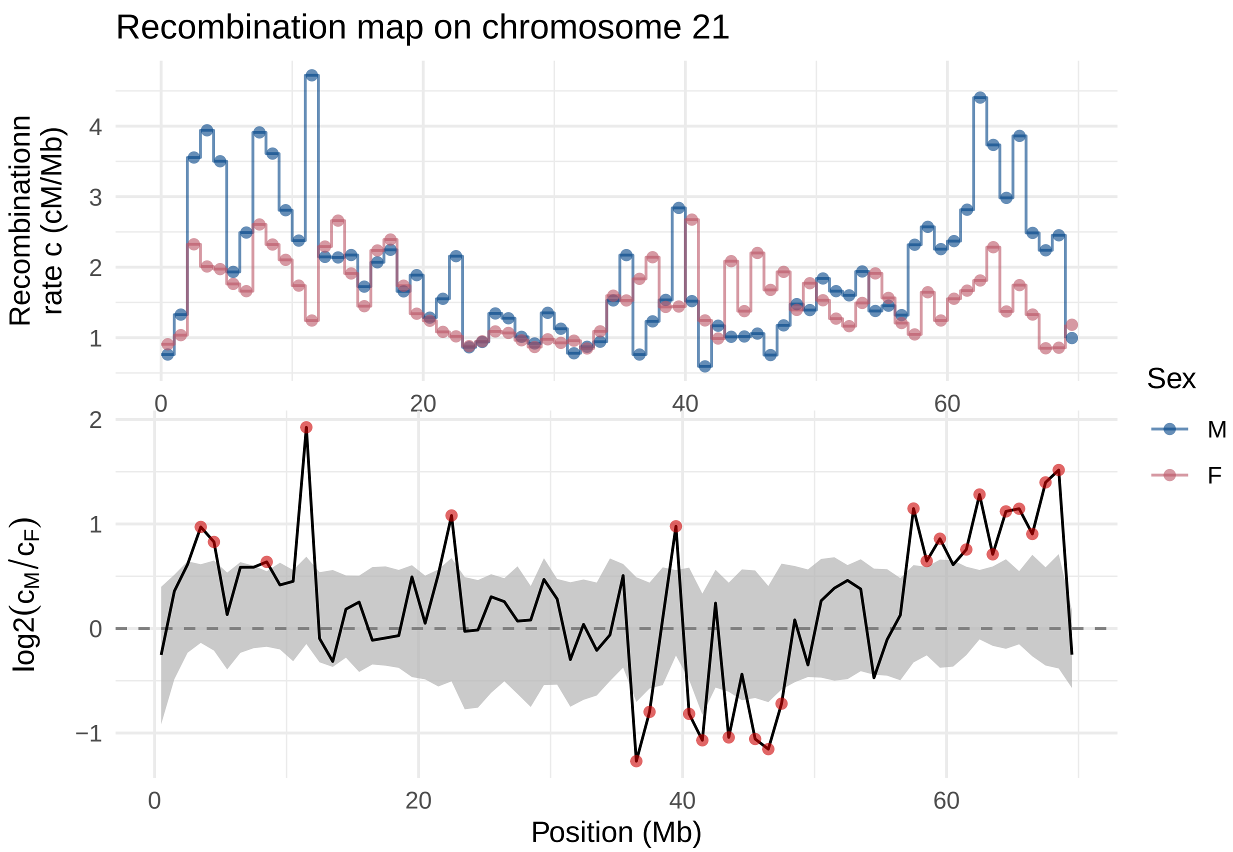
Recombination map on chromosome 21
Recombinationn rate c (cM/Mb)
4
3
2
1
0
20
40
60
Sex
M
F
log2($c_M/c_F$)
2
1
0
−1
0
20
40
60
Position (Mb)

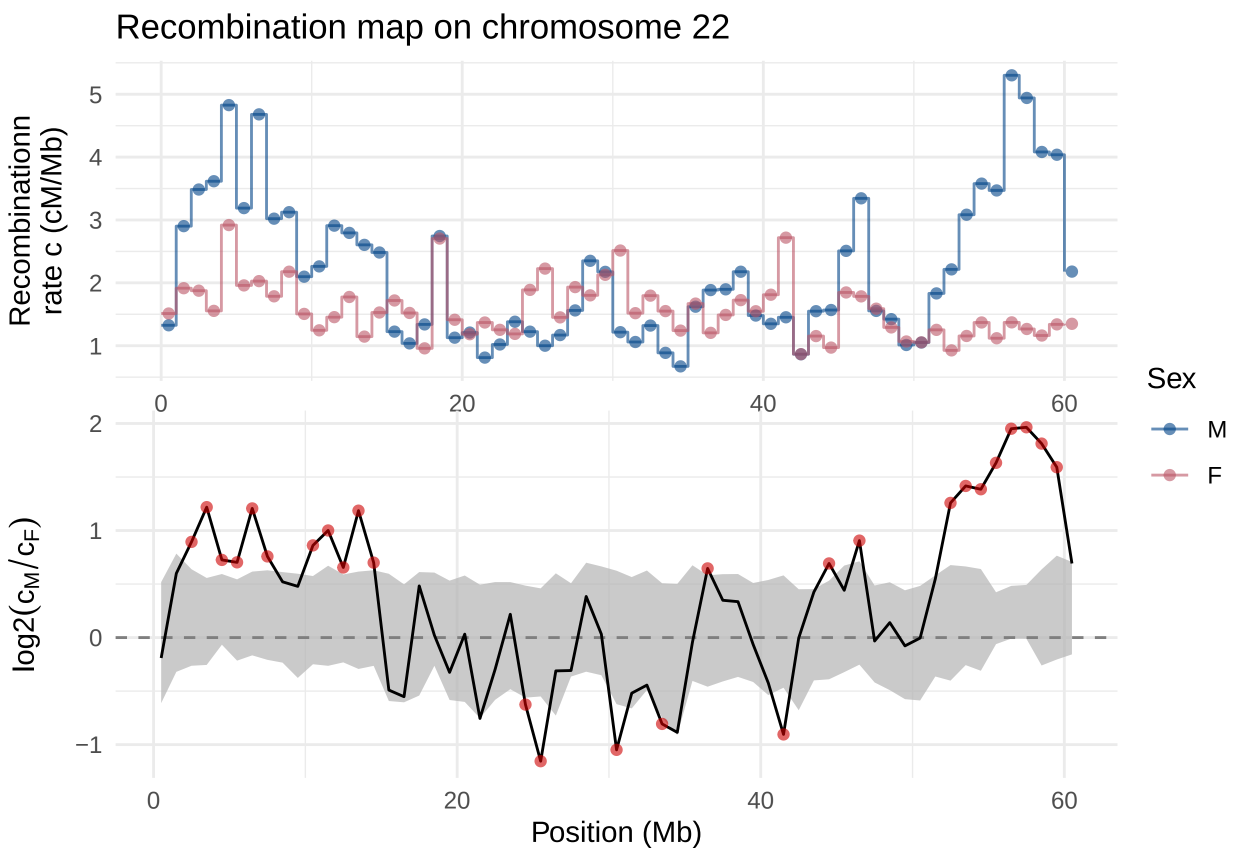

Recombination map on chromosome 22
Recombinationn rate c (cM/Mb)
5
4
3
2
1
0
20
40
60
Sex
M
F
log2(c_M/c_F)
2
1
0
−1
0
20
40
60
Position (Mb)

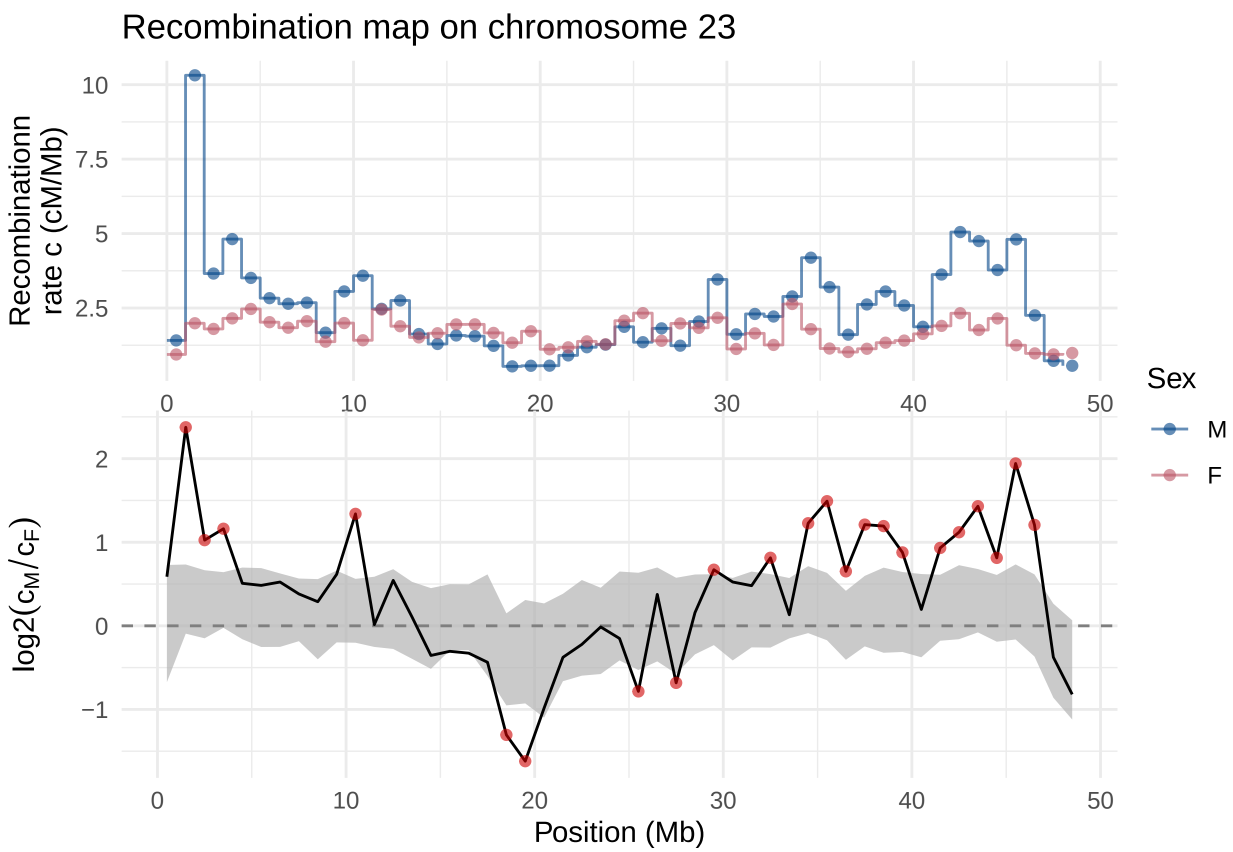
Recombination map on chromosome 23
Recombinationn rate c (cM/Mb)
10
7.5
5
2.5
0
10
20
30
40
50
Sex
M
F
log2(cM/cF)
2
1
0
−1
Position (Mb)

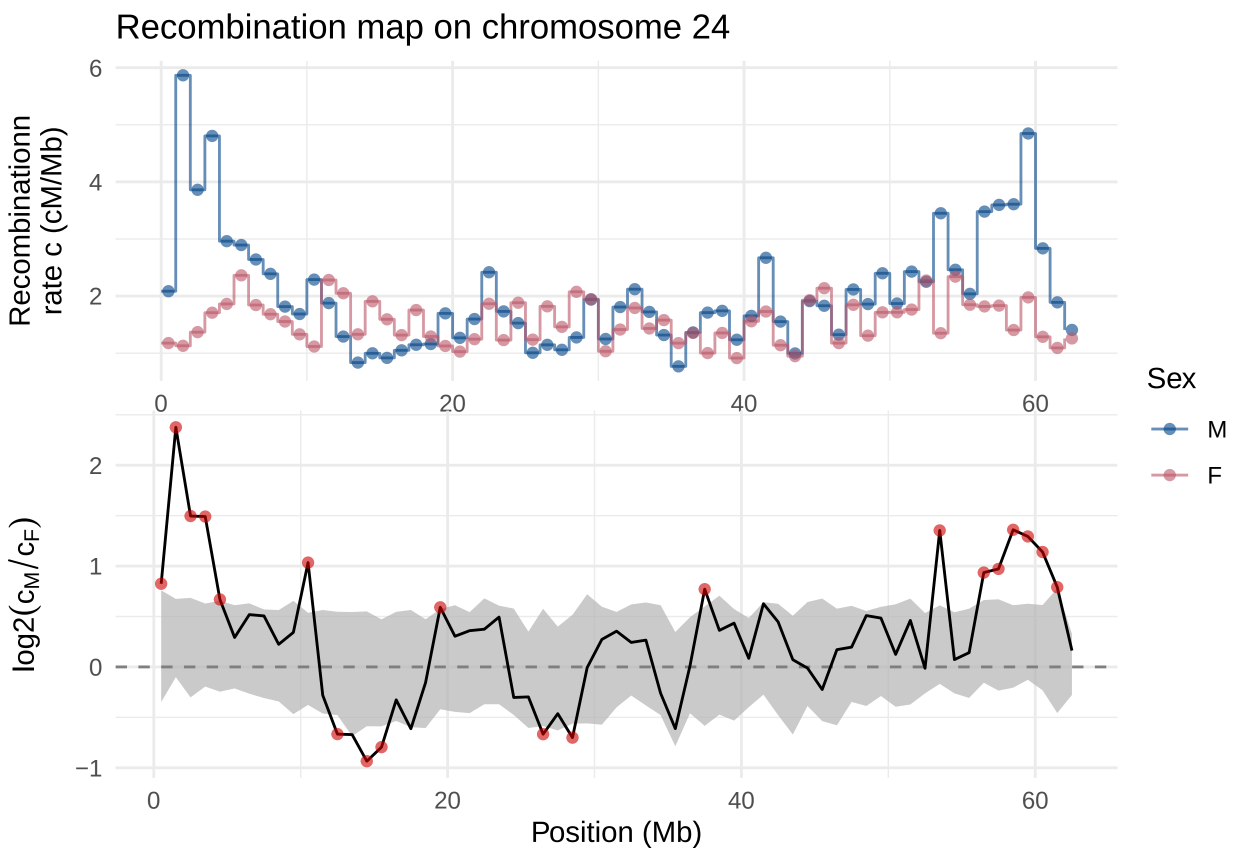
Recombination map on chromosome 24
Recombinationn rate c (cM/Mb)
log2(cM/cF)
Position (Mb)
Sex
M
F

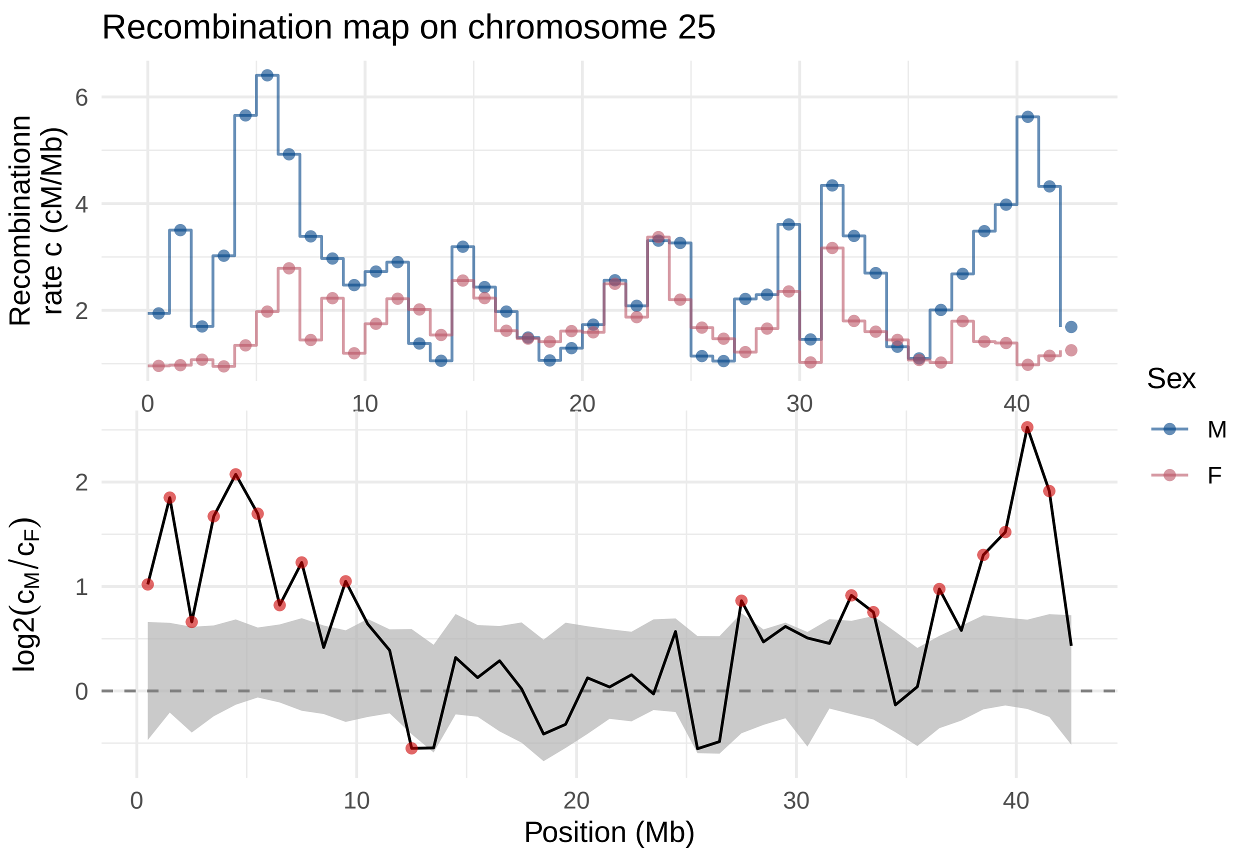

Recombination map on chromosome 25
Recombinationn rate c (cM/Mb)
6
4
2
0
10
20
30
40
Sex
M
F
log2(cM/cF)
2
1
0
0
10
20
30
40
Position (Mb)

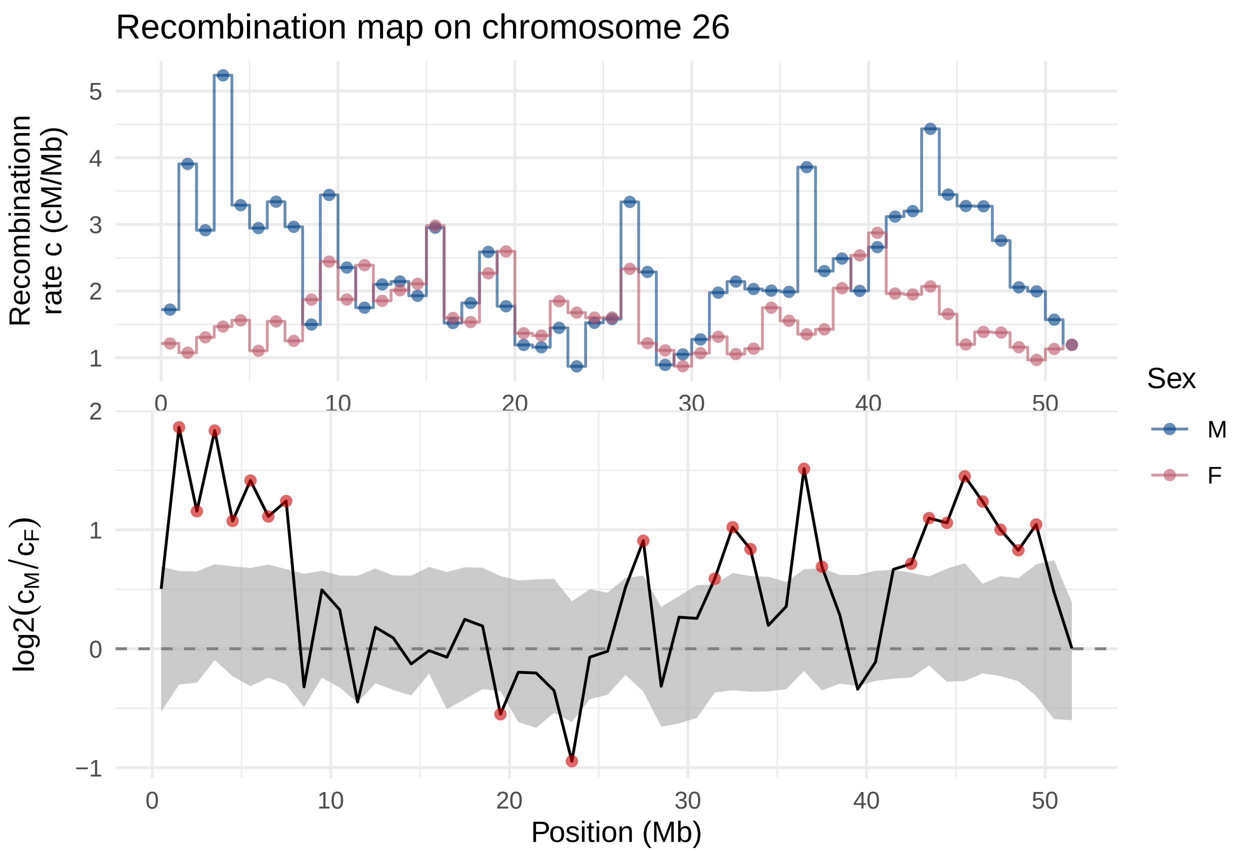

Recombination map on chromosome 26
Recombinationn
rate c (cM/Mb)
log2(cM/cF)
Position (Mb)
Sex
M
F

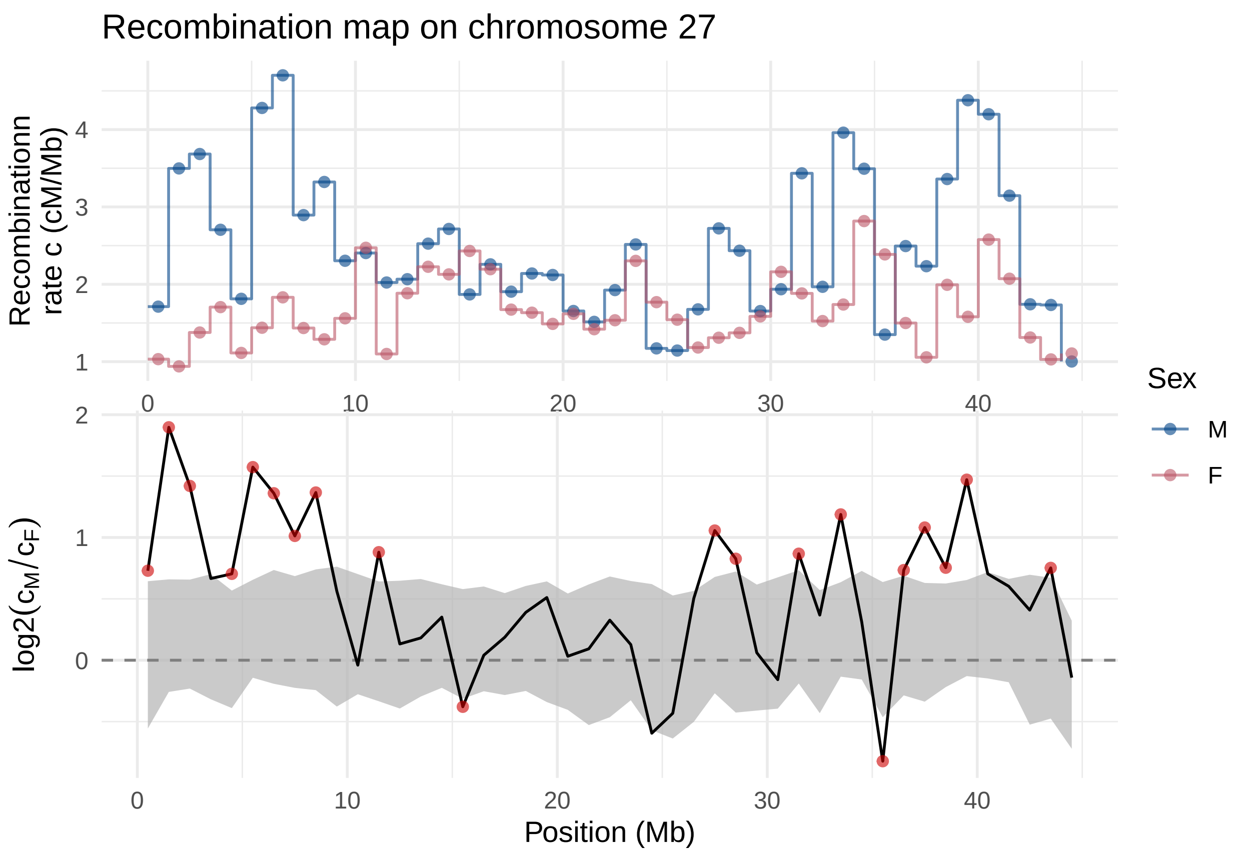
Recombination map on chromosome 27
Recombinationn rate c (cM/Mb)
log2(cM/cF)
Position (Mb)
Sex
M
F
0
10
20
30
40
1
2
3
4
0
1
2

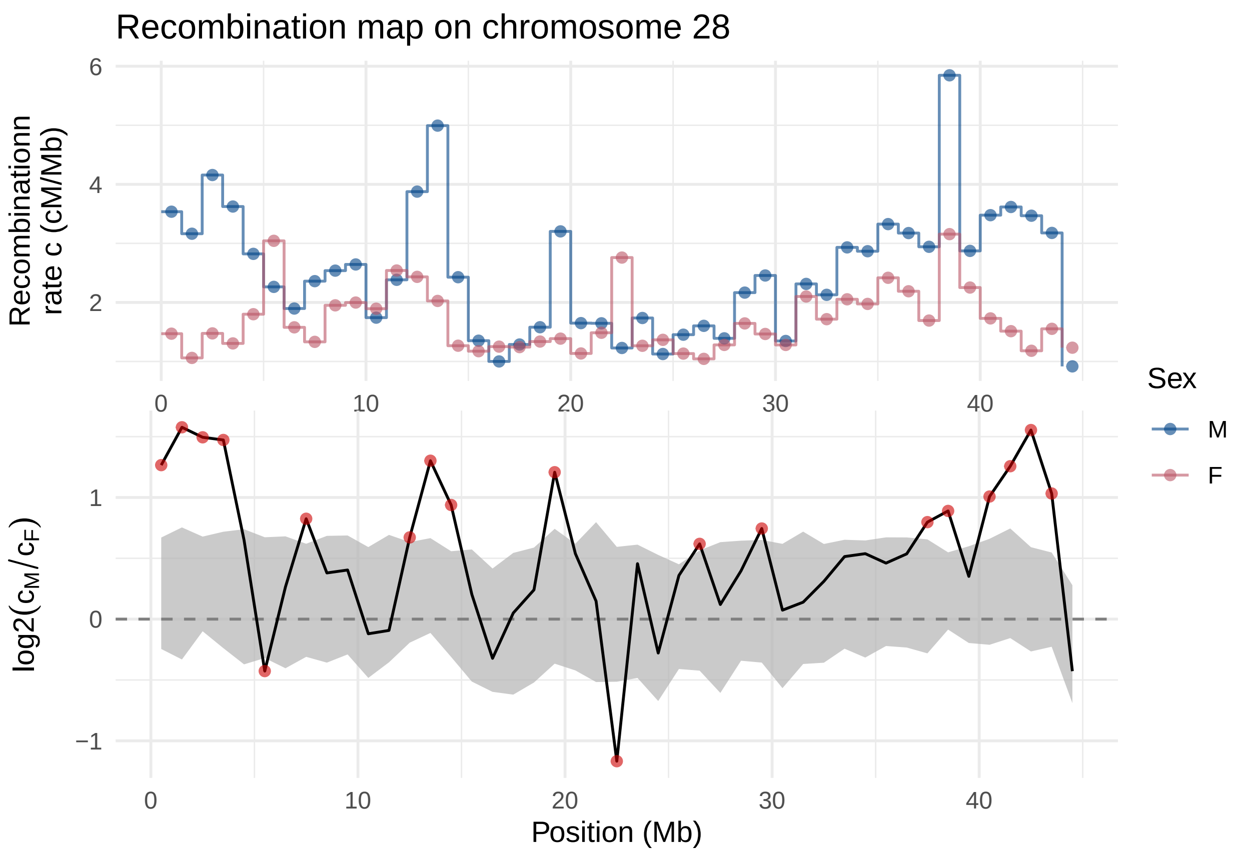
Recombination map on chromosome 28
Recombinationn rate c (cM/Mb)
log2(cM/cF)
Position (Mb)
Sex
M
F

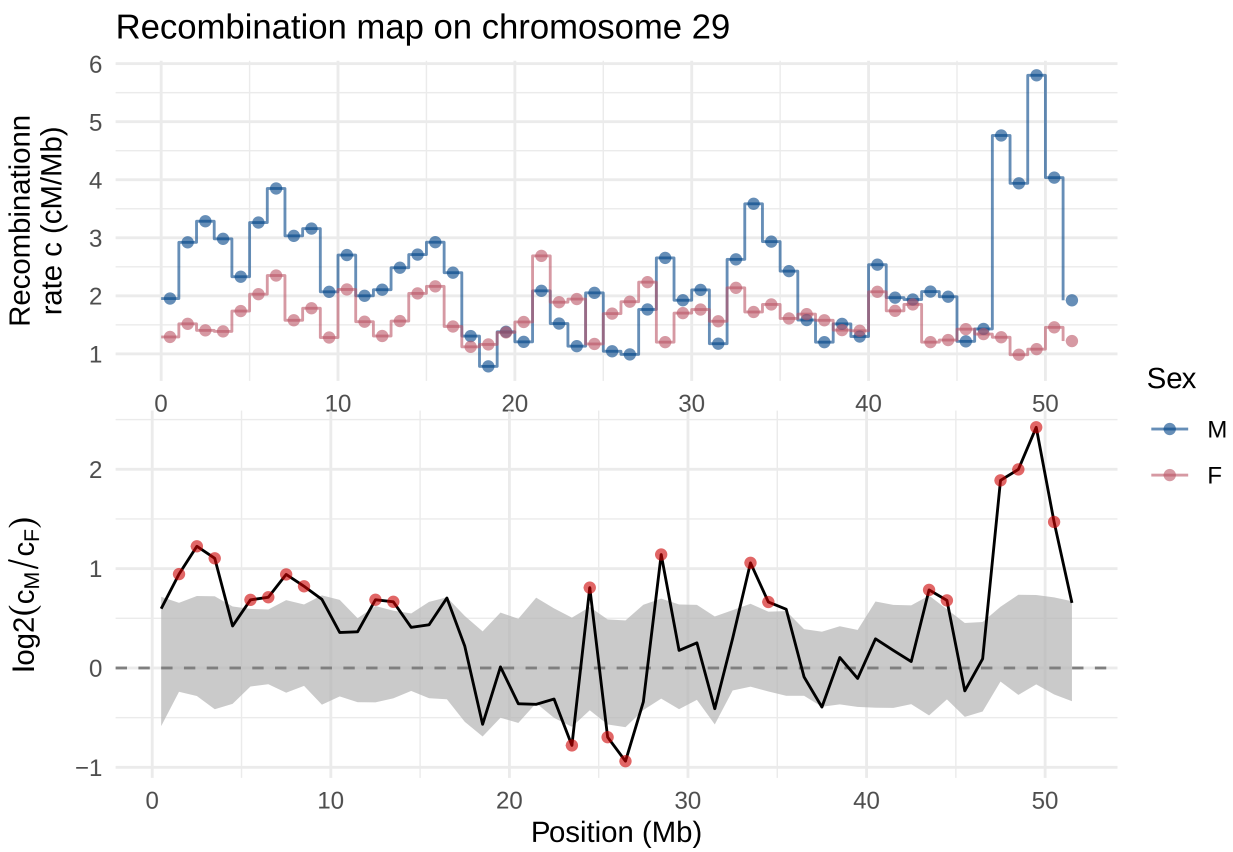
Recombination map on chromosome 29
Recombinationn
rate c (cM/Mb)
6
5
4
3
2
1
0
10
20
30
40
50
Sex
M
F
log2(c_M/c_F)
2
1
0
−1
0
10
20
30
40
50
Position (Mb)
